# CDNF rescues motor neurons in three animal models of ALS by targeting ER stress

**DOI:** 10.1101/2020.05.05.078618

**Authors:** Francesca De Lorenzo, Patrick Lüningschrör, Jinhan Nam, Federica Pilotto, Emilia Galli, Päivi Lindholm, Cora Rüdt von Collenberg, Simon Tii Mungwa, Sibylle Jablonka, Julia Kauder, Susanne Petri, Dan Lindholm, Smita Saxena, Michael Sendtner, Mart Saarma, Merja H. Voutilainen

**Affiliations:** Institute of Biotechnology, HiLIFE, P.O. Box 56, Viikki Biocenter, University of Helsinki, FIN-00014, Helsinki, Finland; Institute of Clinical Neurobiology, University Hospital Würzburg, 97078 Würzburg, Germany; Department of Neurology, Inselspital University Hospital, University of Bern, CH-3010 Bern, Switzerland; Department of Neurology, Hannover Medical School, 30625 Hannover, Germany; Medicum, Department of Biochemistry and Developmental Biology, Faculty of Medicine, University of Helsinki, Helsinki, Finland; Minerva Foundation Institute for Medical Research, Helsinki, Finland

## Abstract

The role of chronic endoplasmic reticulum (ER) stress in the pathophysiology of Amyotrophic lateral sclerosis (ALS), as well as a potential drug target, has received increasing attention. Here, we investigated the mode of action and therapeutic effect of the ER resident protein cerebral dopamine neurotrophic factor (CDNF) in preclinical models of ALS harboring different genetic mutations. We identify that intracerebroventricular (i.c.v.) administration of CDNF significantly halts the progression of the disease and improves motor behavior in TDP43-M337V and SOD1-G93A rodent models of ALS. CDNF rescues motor neurons (MNs) *in vitro* and *in vivo* from ER stress associated cell death and its beneficial effect is independent of genetic disease etiology. Notably, CDNF regulates the unfolded protein response (UPR) initiated by transducers IRE1α, PERK, and ATF6, thereby enhancing MN survival. Thus, CDNF holds great promise for the design of new rational treatments for ALS.

## INTRODUCTION

Amyotrophic lateral sclerosis (ALS) is a progressive motor disorder characterized by the dysfunction and death of motor neurons (MNs) in the spinal cord, brainstem, and motor cortex, which results in the atrophy of skeletal muscles and paralysis. Any cure or disease-modifying therapy is currently lacking. ALS patients die within 3 to 5 years of diagnosis and respiratory failure is the most common cause of death ^1^. To date, ALS etiology remains mostly unknown: only about 5-10% of the cases are familial, while the remaining 90-95% of the cases occur sporadically, indicating the influence of multiple factors in ALS pathogenesis. Accumulation of misfolded and aggregated proteins leads to endoplasmic reticulum (ER) stress, which eventually activates the unfolded protein response (UPR).

The UPR is a physiological signaling cascade that suppresses protein translation, degrades misfolded proteins and facilitates protein folding. The UPR consists of three pathways, initiated by transmembrane ER sensors at the ER membrane: inositol-requiring enzyme 1 (IRE-1), protein kinase RNA-like endoplasmic reticulum kinase (PERK), and activating transcription factor 6 (ATF6). At onset, the UPR is protective, but, under prolonged ER stress conditions, the UPR promotes apoptotic pathways ^2^.

Substantial evidence supports the involvement of chronic ER stress in the pathophysiology of MN degeneration in ALS, both in patients and animal models. UPR markers were found to be upregulated in the spinal cord of sporadic and familial ALS patients ^3–6^ and increased amount of protein disulfide isomerase (PDI) was detected in the cerebrospinal fluid of sporadic patients ^3^. Transcriptional analysis of MNs derived from iPSCs of ALS patients revealed that UPR alterations were conserved among MNs harbouring SOD1 or C9ORF72 mutations, emphasizing the susceptibility of MNs to ER stress ^7–9^.

Fast-fatigable MNs in the SOD1 transgenic mice were shown to be intrinsically more sensitive to ER stress ^10^ and an analogous upregulation of UPR markers was also reported in other studies on this mouse model ^11,12^. Interestingly, a recent work revealed that, under chronic ER stress conditions, also wild type (WT) SOD1 protein tends to aggregate in mice, forming abnormal species ^13^. Similarly, pharmacological induction of ER stress causes TDP43 to accumulate and, *vice versa*, overexpression of mutated TDP43 triggers ER stress ^14^. Genetic ablation of the ER chaperone SIL-1 in the SOD1-G93A mouse model enhanced ER stress and consequently exacerbated the disease progression, emphasizing the importance of ER homeostasis in ALS pathophysiology ^15^. Furthermore, translocation of FUS protein from the nucleus to the cytoplasm has been linked to ER stress ^16,17^. Additionally, mutations in genes encoding for ER proteins have been linked to familial and sporadic cases of ALS. A missense mutation E102Q in the ER chaperone Sigma-1 receptor gene has been reported in few familial ALS cases ^18^. A total of nine PDIA1 missense variants and seven PDIA3 missense variants for the gene encoding PDI were identified in sporadic ALS patients ^19^.

Compounds interfering specifically with the PERK/p-eIF2alpha pathway or the knockdown of its downstream effector ATF4 partially alleviate ALS symptoms in the SOD1-G93A rodent model, providing neuroprotection and delaying disease progression ^10,20–22^. Inhibiting the IRE1α -XBP1 arm of UPR signalling since embryonic stage revealed a beneficial role in the disease pathophysiology, due to an enhanced clearance of mutant SOD1 aggregates by autophagy pathways ^4^. Moreover, targeting ER folding capacity and homeostasis by overexpression of SIL-1 in the SOD1-G93A mouse model proved to be neuroprotective and prolonged survival ^15^.

Until now, only inhibitors of individual UPR pathways have been described, but due to the double-edged sword nature of UPR, molecules modulating all three pathways are needed. Despite efforts to find drugs targeting ER stress to treat protein-misfolding and aggregation disorders, its value as a pharmacological target warrants further exploration.

Cerebral dopamine neurotrophic factor (CDNF) is an ER-resident protein expressed in the central nervous system mostly in neurons and peripheral tissues, including skeletal muscles, the target tissue of MNs ^23^. By virtue of its C-terminal signal KTEL, which resembles the canonical Lys-Asp-Glu-Leu (KDEL) sequence for ER retention, CDNF-protein is primarily located in the ER lumen and secretion of mouse CDNF is regulated by the ER-resident proteins GRP78 and KDEL-R1 ^24^, hinting at a possible involvement in ER homeostasis. When overexpressed or microinjected into neurons, CDNF rescues cells from ER stress-induced apoptosis. Most importantly, CDNF protects and restores dopamine neurons in rodent and non-human primate models of Parkinson’s disease (PD) ^23,25–27^. CDNF was tested in Phase I-II trial on Parkinson’s patients and the study met primary endpoints of safety and tolerability at 12 months (Clinical trial number NCT03295786) ^28^.

In order to determine the efficacy and potential of CDNF as a drug candidate for ALS, we treated three independent rodent models of ALS: (i) a severe fast progressing TDP43-M337V rat model, previously described by Huang *et al.* ^29^; (ii) the well characterized SOD1-G93A mouse model; (iii) a novel pre-symptomatic TDP43-M337V mouse model. Albeit being very different from each other, all three models exhibited signs of ER stress and death of MNs. Our results reveal that CDNF markedly modulates disease symptoms and rescues MNs from ER stress-induced cell death, indicating that ER stress is an important component of the pathophysiology of this disease, irrespective of its genetic etiology. Moreover, our data provide evidence that CDNF mediated inhibition of ER stress at symptomatic stages has the capacity as a therapeutic intervention for this disorder.

## RESULTS

### Treatment with CDNF rescues MNs and improves motor behavior in a TDP43-M337V rat model

To investigate the potential neuroprotective effect of CDNF in ALS, we started administering CDNF in the ChAT-tTA/TRE-hTDP43-M337V rat model introduced by Huang *et al.* ^29^. In this preclinical model, the expression of human mutated TDP43 protein was dependent on the choline acetyl-transferase (ChAT) promoter via a Tet-regulatory system. The transgene was kept inactive until adult age by administration of doxycycline (Dox) in drinking water (50 mg/ml). At the age of 60 days, Dox was completely withdrawn from the water and expression of TDP43-M337V protein became evident in the spinal cord MNs of the rats already three days later. In the original study, the model showed a severe disease development, as the rats exhibited signs of motor impairment within a week and reached paralysis in one week after symptoms onset. At this stage, over 60% of spinal MNs were lost in the transgenic rats ^29^. As the progression of the disease was very fast and, therefore, the therapeutic window was narrow, we decided to implement the protocol and opted for a partial withdrawal of Dox, instead of a complete one. Starting from 60 days of age, rats were administrated Dox in drinking water at a concentration of 10 mg/ml. The partial withdrawal of Dox induced a slower progression of the disease (**Fig. 1b**). A continuous infusion of 6 μg/day of CDNF or phosphate buffered saline (PBS) as vehicle in the brain lateral ventricle of TDP43-M337V and WT littermates was achieved by using Alzet minipumps. The minipumps, connected through a catheter to a cannula directly infusing in the cerebral ventricle (i.c.v.), were implanted one week before the activation of the transgene and continued to administer CDNF or the vehicle for 28 days. After the partial withdrawal of Dox, the rats were monitored for weight and motor behavioral changes for three weeks (**Fig. 1a**). Upon transgene activation, no difference in the rotarod performance was detected between the TDP43-M337V rats and the WT littermates until day 10. The ability of running on the rotating rod decreased substantially in the transgenic rats from day 10 onwards and major signs of hind limb stiffness and gait impairment were detected by day 20 (**Fig. 1b**). A continuous i.c.v. administration of CDNF significantly ameliorated the motor performance of treated TDP43-M337V, which was comparable to WT littermates. No difference over time in the latency to fall was detected in WT rats (**Fig. 1b**). Moreover, we verified whether the improvement in the motor performance of CDNF treated rats compared to PBS ones correlated with the number of MNs in the lumbar spinal cord. Indeed, we found that the transgenic rats that received PBS lost about 50% of the lumbar spinal MNs but treatment with CDNF significantly increased the number of surviving MNs in the same area to about 75% in comparison to the WT control (**Fig. 1c-d**).

**Figure 1.**
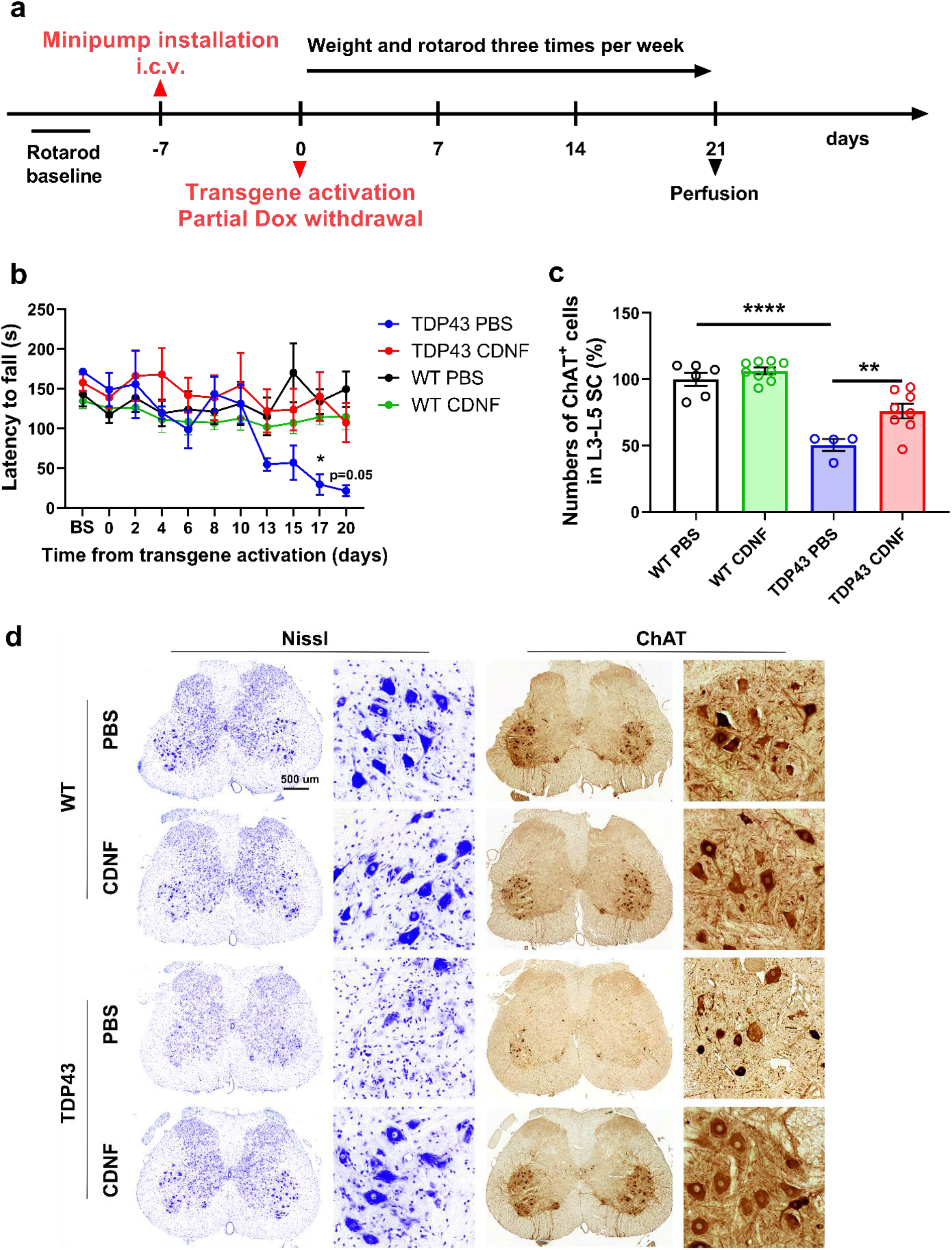
Continuous i.c.v infusion of CDNF improves motor behavior and protects spinal MNs in the ChAT-tTA/TRE-TDP43-M337V rat model. (**a**) Experimental design: upon reaching adult age, Alzet minipumps connected to a catheter were implanted in ChAT-tTA/TRE-TDP43-M337V and WT rats to infuse CDNF (6 μg/day) or PBS. One week later, the activation of transgene was induced by partial withdrawal of Dox. Rats were monitored for weight and motor behavior changes until day 21 from transgene induction, when the rats were perfused. (**b**) Latency to fall of PBS/CDNF-treated transgenic and WT littermates recorded three times per week. BL=baseline. (**c**) Quantification and comparison of the number of MNs in the lumbar (L) spinal cord, area L3-L5. (**d**) Representative images of Nissl and ChAT^+^ MNs in the lumbar spinal cords. Scale bar 500 μm. Mean ± SEM, n=4-9/group in **b-d**. *p<0.05, **p<0.01, ****p<0.0001, repeated measures ANOVA followed by Tukey post-hoc test in **b** and one-way ANOVA followed by Tukey post-hoc test in **c**.

### CDNF attenuates the ER stress response in the TDP43-M337V *in vitro* and *in vivo*

Next, we sought to identify whether the protective effect of CDNF lay in its ability to reduce the ER stress response, as previous evidence has hinted to a possible role of CDNF in the ER homeostasis ^24,30^. Thus, we examined the effect of CDNF on TDP43 expressing MNs *in vitro*, isolated from embryonic day 13 (E13) mice. WT TDP43 and mutated TDP43-M337V expressing neurons were pharmacologically stressed with thapsigargin (TP) or tunicamycin (TM), to induce ER stress, and then treated with CDNF. The levels of phosphorylated eIF2α (p-eIF2α) and CHOP (gene *Ddit3*) were reduced by CDNF treatment (**Fig. 2a-d**), indicating that CDNF attenuates ER stress in this model. The levels of phosphorylated TDP43 and the total levels of the protein did not change upon CDNF treatment (**Fig. S1**). To further investigate the neuroprotective mechanism behind CDNF action in the ChAT-tTA/TRE-hTDP43-M337V rat model, we sacrificed the rats 21 days after the transgene activation and collected the spinal cord for immunohistochemical analyses (**Fig. 1a**). We found that the levels of ER chaperone GRP78 (*alias* BiP, gene *Hspa5*) and phosphor-PERK (p-PERK) were highly upregulated in the vehicle-treated TDP43-M337V rats compared to WT controls (**Fig. 3a-f**). This increase in the ER stress response in this model was not previously described in the literature and strengthens the hypothesis of the importance of ER stress in the pathophysiology of ALS. Upon treatment with CDNF, the expression of p-PERK and GRP78 was significantly reduced in the ventral horn of the lumbar spinal cord and, specifically, in the ChAT^+^ MNs (**Fig. 3a-f**).

**Figure 2.**
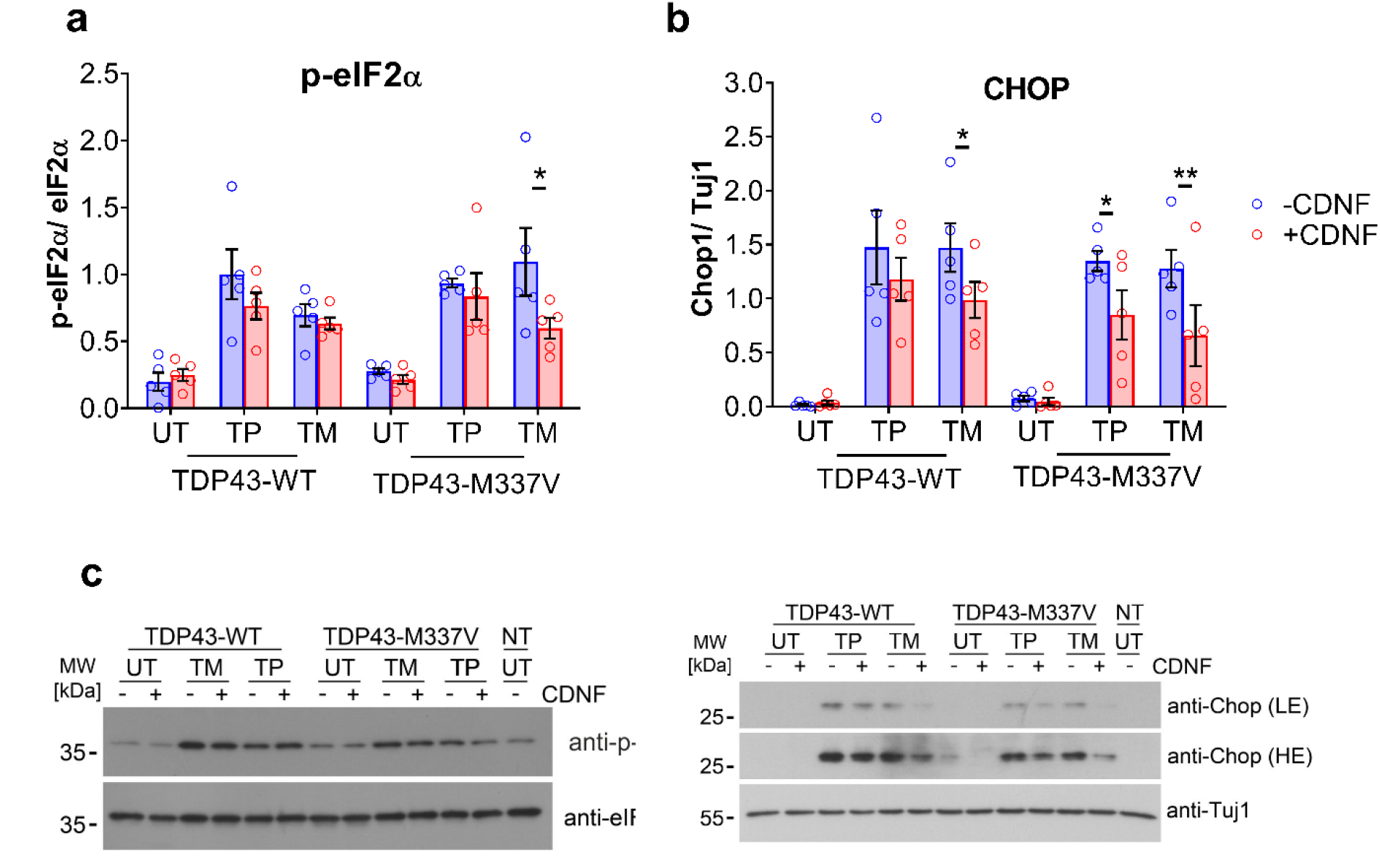
CDNF treatment decreases UPR markers expression in TDP43-M337V expressing MNs. (**a-b**) Protein expression of UPR markers phosphorylated eIF2α and CHOP in the MNs expressing TDP43 WT and TDP43-M337V, which were previously stressed with thapsigargin (TP) and tunicamycin (TM). (**c-d**) Representative blots of p-eIF2 α and CHOP. Mean ± SEM of 5 different experiments in **a-b**. *p<0.05, **p<0.01; two way ANOVA followed by Sidak post-hoc test in **a-b**.

**Figure 3.**
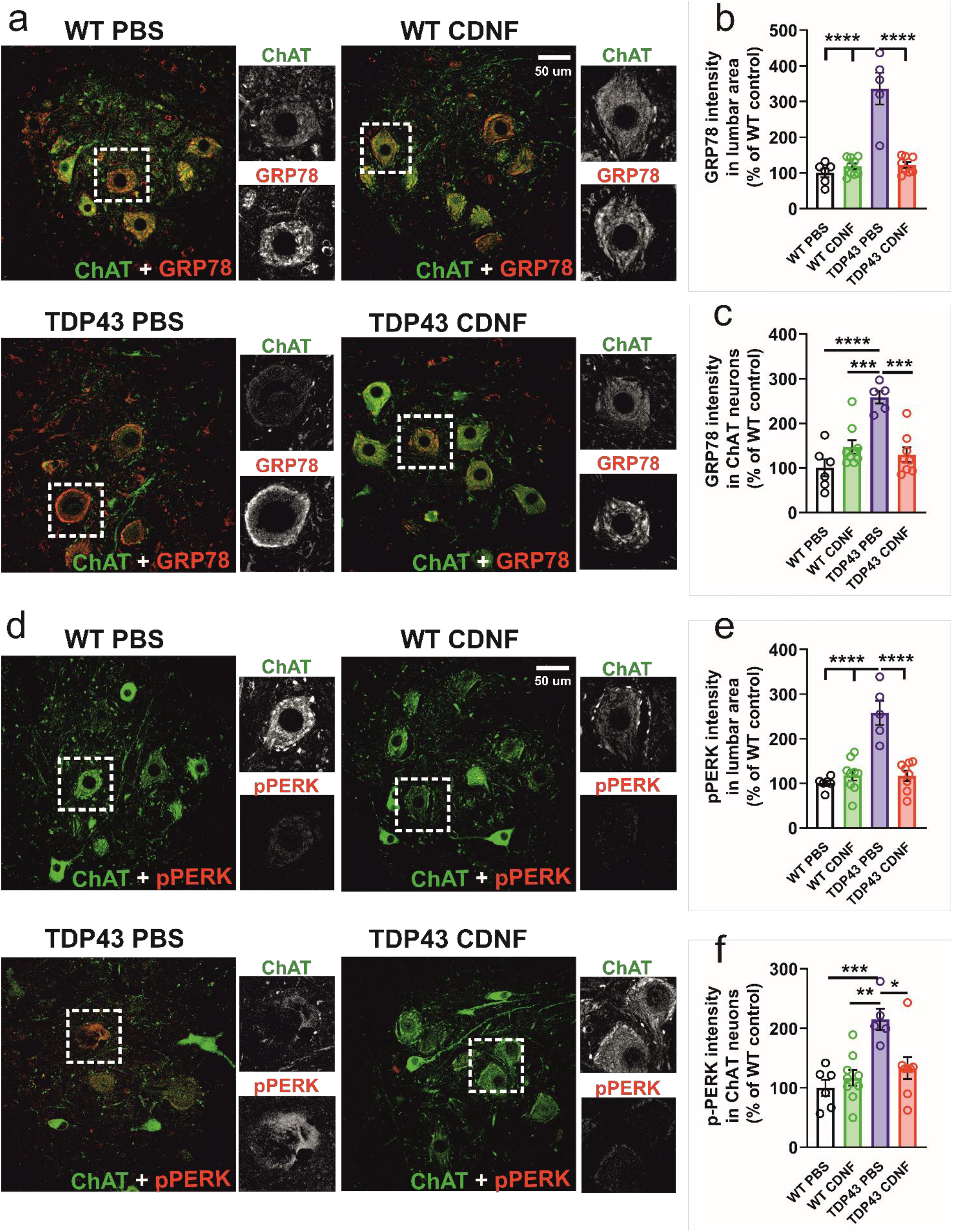
CDNF administration attenuates the expression of UPR markers in the spinal cord of ChAT-tTA/TRE-TDP43-M337V rats at 21 days after transgene activation. (**a**) Representative fluorescence images of ChAT (Green) and GRP78 (Red) protein expression in PBS/CDNF-treated ChAT-tTA/TRE-TDP43-M337V compared to PBS/CDNF-treated WT rats. Scale bar 50 μm. (**b**) Quantification of the pixel value of GRP78 in the lumbar ventral horn area. (**c**) Quantification of the pixel value of GRP78 in the ChAT^+^ MNs. (**d**) Representative fluorescence images of ChAT (Green) and p-PERK (Red) protein expression in PBS/CDNF-treated ChAT-tTA/TRE-TDP43-M337V compared to PBS/CDNF-treated WT rats. Scale bar 50 μm. (**e**) Quantification the of pixel value of p-PERK in the lumbar ventral horn area. (**f**) Quantification of pixel value of pPERK within ChAT^+^ MNs. Mean ± SEM, n=5-9/group. *p<0.05; **p<0.01; ***p<0.001; ****p<0.0001 one-way ANOVA followed by Tukey post-hoc test in **b**, **c**, **e**, and **f**.

### A single i.c.v. injection of CDNF ameliorates motor behavior, improves survival, and rescues lumbar spinal MNs in the SOD1-G93A mouse model

To determine whether the effect of CDNF would be specific only to the TDP43 model or if it could be applied to any model affected by ER stress, we treated SOD1-G93A mice with CDNF. In this well characterized ALS model, chronic ER stress response in the corresponding MNs has been previously reported ^10,11^. To this end, we utilized a single injection of CDNF, which in PD animal models effectively counteracted dopamine neuron degeneration ^23^. As proof-of principle, we first confirmed that CDNF and ^125^I-labelled CDNF injected in the brain lateral ventricle efficiently diffuse to different areas of the brain, including cortex, striatum, and substantia nigra, and to the lumbar spinal cord (**Fig. S2**). Moreover, we found that CDNF specifically co-localizes with lumbar MNs (**Fig. S3**). Early symptomatic SOD1-G93A mice and WT littermates of 13 weeks of age received a single i.c.v. injection of 10 μg of human CDNF or phosphate buffered saline (PBS) as vehicle and were examined twice a week to follow changes in symptoms and motor behavior until the final stage of paralysis (**Fig. 4a**). At the time of treatment, SOD1-G93A mice displayed measurable tremors in the hind limbs. Upon a single CDNF injection, SOD1-G93A mice developed gait impairment and paralysis symptoms significantly more slowly than PBS-treated mutant littermates (**Fig. S4a-b**), and survival was increased in both female (7.1%) and male (7.6%) SOD1 mice (**Fig. 4b-c**). Along with this, their balance and motor behavior performance were significantly ameliorated by CDNF treatment. In the accelerating rotarod, CDNF-treated female and male mice showed an increased latency to fall compared to vehicle treated-mice. No statistical differences were found in CDNF or vehicle-injected WT littermates over time (**Fig. 4d-e**). One week after CDNF or PBS injection, the CDNF group showed an increased ability to run on the smallest 8 mm rod in the multiple static rods experimental paradigm for females and on the 21 and 11 mm rods for males (**Fig. 4f-g**), compared to the PBS group. These gender differences in the experimental tasks are in accordance with previous observations that males develop major symptoms approximately one week earlier than female littermates ^31,32^. In the open field, SOD1-G93A mice exhibited a decreased number of rearings compared to WT mice, which is associated with less strength in the hind limb muscles, and CDNF treatment increased the number of rearings compared to PBS controls at 16 weeks (**Fig. 4h**). Moreover, immunohistochemical analyses of the lumbar spinal cord revealed a significantly higher number of MNs present in the CDNF-treated compared to PBS-treated mice, which correlated with the behavioral improvement (**Fig. 4i**).

**Figure 4.**
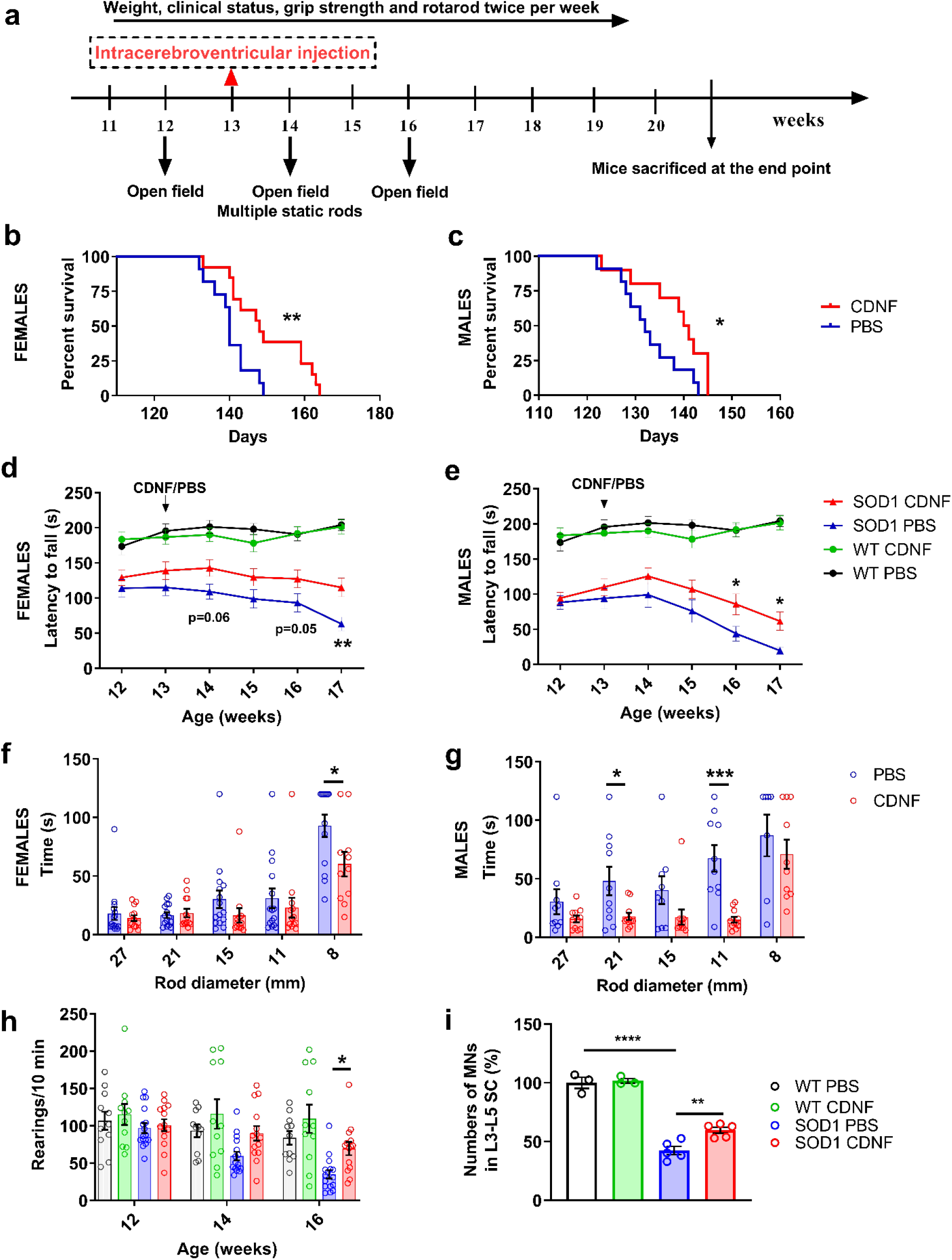
A single i.c.v. injection of CDNF halts symptom progression, ameliorates motor behavior, and improves the survival of mice and spinal MNs in the SOD1-G93A mouse model. (**a**) Early symptomatic SOD1 mice and WT littermates were injected with 10 ug of CDNF or vehicle at 13 weeks of age. Animals were monitored twice per week for weight changes, symptom development, and motor performance. (**b-c**) Survival of SOD1-G93A mice upon a single injection of CDNF or PBS treatment. (**d-e**) Latency to fall of CDNF/PBS treated-SOD1-G93A and WT littermates measured every week. (**f-g**) Travel time of CDNF or PBS-treated SOD1-G93A mice along suspended rods of decreasing diameters (27 to 8 mm) one week after treatment. (**h**) Number of rearings of CDNF/PBS treated-SOD1-G93A and WT littermates in the open field paradigm measured at 1 and 3 weeks after injection. (**i**) Quantification and comparison of the number of MNs in the lumbar spinal cord after CDNF or vehicle treatment. Mean ± SEM, n=10-12/group in **b-c**, n=18-20 for females, n=14-15 for males, n=20 for WT in **d-g**, n=11/group in **h**, n=3 for WT and n=5 for SOD1 in **i**. *p<0.05, **p<0.01, ***p<0.001, ****p<0.0001, log-rank test in **b-c**, repeated measures ANOVA followed by Tukey post-hoc test in **d-e** and **h**; unpaired t-test in **f-g;** one-way ANOVA followed by Tukey post-hoc in **i**.

### CDNF increases survival of SOD1-G93A embryonic MNs after toxin treatment by mitigating all three UPR pathways

We further investigated whether CDNF modulates ER stress and survival of SOD1-G93A-derived embryonic MNs *in vitro* and, in particular, whether its effect is limited to only one branch of the UPR or is extended to all three. MNs from E13 WT and SOD1-G93A mouse embryos, which depend on trophic support for their survival *in vitro*, were cultured either in the presence of established neurotrophic factors, brain-derived neurotrophic factor (BDNF) ^33^ and ciliary neurotrophic factor (CNTF) ^34^ or only with CDNF. Interestingly, CDNF alone had a significant effect on the survival of the MNs as compared to the untreated conditions, though not to the same degree as observed with the combination of BDNF and CNTF (**Fig. 5a**). No difference in the morphology of cultured MNs was observed between any of the aforementioned culture conditions (**Fig. S5**). SOD1-G93A mutant MNs were obtained from a mouse model of ALS, which develops first clinical symptoms at about three months after birth. However, when ER stress was induced in cultured MNs by TP treatment, these embryonic SOD1-G93A mutant MNs appeared significantly more sensitive compared to MNs from WT littermates, supporting previous observations made with this mouse model ^10^. This sensitivity was seen despite the continuous presence of BDNF and CNTF in culture, but when CDNF was added to ER stressed SOD1-G93A mutant MNs, survival was rescued to the levels observed in WT MNs (**Fig. 5b**). These experiments indicate that CDNF differs from established neurotrophic factors by its specific activity in counteracting ER stress-mediated cell death, while it has little effect on non-stressed cells. The protective effect of CDNF in SOD1-G93A MNs from TP-induced ER-stress occurred in a dose-dependent manner and at a concentration of 20 ng/ml or higher the survival of SOD1-G93A MNs was comparable to that of WT cells (**Fig. 5c**). Next, we analyzed whether CDNF, when compared to BDNF/CNTF, can modulate the signaling pathways of the UPR in cultured MNs treated with either TP or TM. We observed a partial translocation of ATF6 protein from the cytoplasm to the nucleus of the neurons upon treatment with TP or TM. However, the addition of CDNF significantly reduced ATF6 translocation in SOD1-G93A treated with both TP or TM (**Fig. 5d-e**). Furthermore, CDNF diminished the splicing levels of *xbp1s* transcripts observed after TP or TM treatment, particularly in SOD1-G93A MNs, suggesting an inhibition of the IRE-1α-linked pathway (**Fig. 5f**). Additionally, CDNF treatment significantly decreased p-eIF2α protein levels and CHOP, activated downstream of the PERK signaling pathway, in TP and in TM treated SOD1-G93A MNs (**Fig. 5g-i, S6**). Collectively, these results show that SOD1-G93A MNs are already at embryonic stage intrinsically more sensitive to ER stress and that CDNF efficiently counteracts the ER stress-induced cell death by reducing the activity of all three major UPR signaling pathways. This effect is in contrast to that of the available inhibitors of ER stress targeting only one of the three pathways.

**Figure 5.**
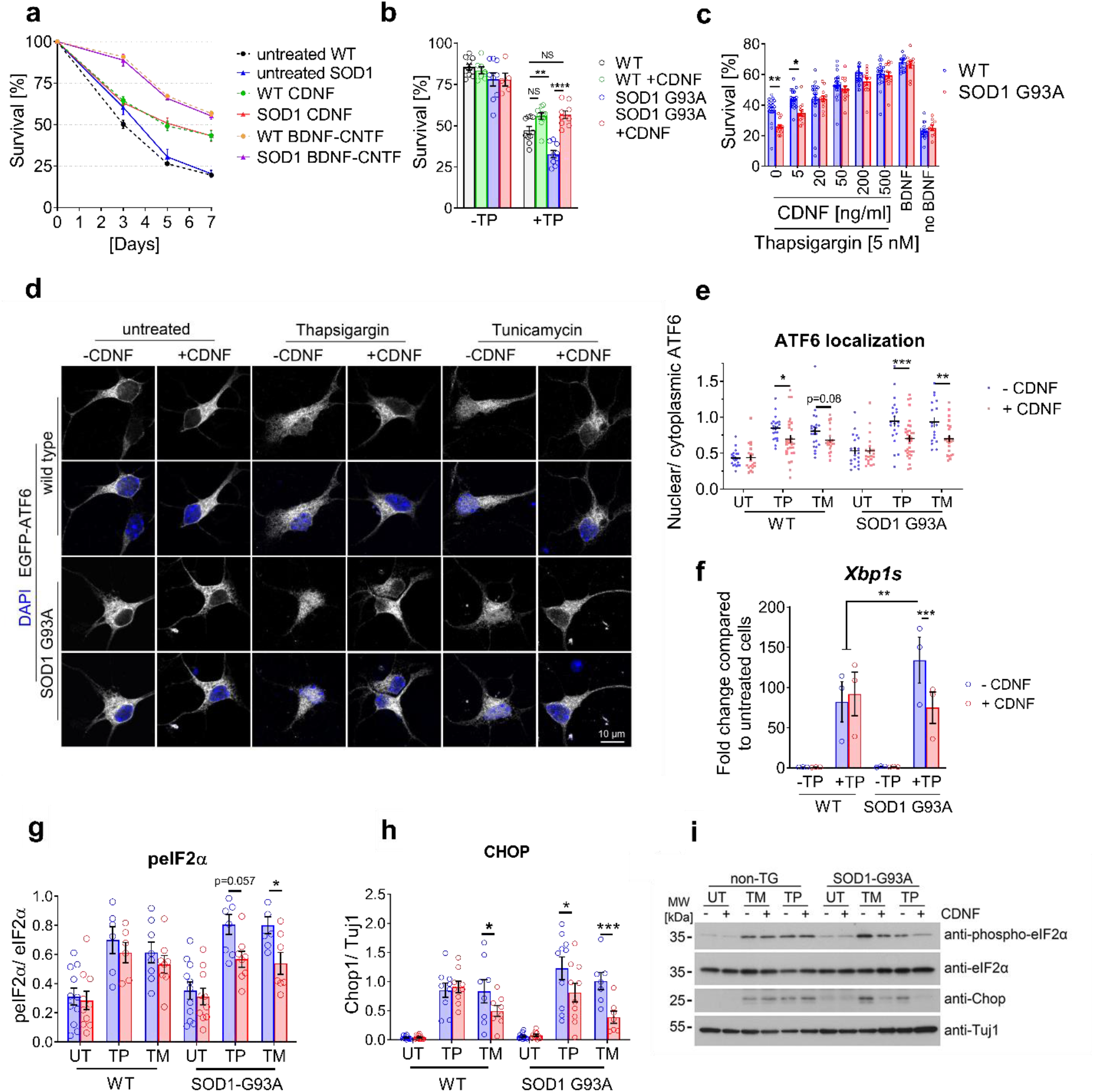
CDNF protein administration increases the survival of SOD1-G93A mouse-derived embryonic MNs and reduces the expression of UPR markers after toxin treatment. (**a**) Survival of WT and SOD1-G93A embryonic MNs upon CDNF or CNTF and BDNF treatment. (**b**) Effect of CDNF administration on WT and SOD1 MNs survival upon 5 nM thapsigargin (TP) treatment or no treatment. (**c**) Dose-response curve of CDNF administration upon TP treatment on WT and SOD1-G93A MNs survival. (**d**) Effect of CDNF on ATF6 translocation from cytoplasm to nucleus in MNs treated with TP or tunicamycin (TM). (**e**) Quantification of **d**. (**f**) mRNA expression of UPR markers *Xbp1s* in the MNs treated as in Figure **b**. (**g-h**) Expression levels of eIF2α phosphorylation and CHOP protein, respectively, in the MNs treated as in Figure **d**. (**i**) Representative blots for the quantification in **g** and **h**. Mean ± SEM of 3 independent experiments in **a**, **b** and **f**, 4 experiments in **c-e** and 7 experiments in **g-i** *p<0.05; **p<0.01; ***p<0.001, two-way ANOVA followed by Bonferroni post-hoc test in **b** and **f**, and Sidak’s post-hoc test in **c, e, g** and **h**. TG=transgenic.

### CDNF attenuates all three UPR branches initiated by PERK, IRE-1α and ATF6 *in vivo*

To investigate the mechanisms involved in the *in vivo* action of CDNF in the SOD1-G93A model, we analyzed the UPR signaling pathways in lumbar MNs isolated via laser microdissection (**Fig. 6a**) at 13 and 17 weeks, i.e. 5 days after CDNF i.c.v. injections (**Fig. 6b-f**). Interestingly, at 13 weeks we only found an upregulation of *Atf4* and *Chop* transcripts in SOD1-G93A mice. As both molecules are downstream effectors of the PERK pathway, which blocks general protein translation initiation in stressed cells, these data suggest that this branch of UPR signaling is the first to be activated in lumbar MNs of SOD1-G93A mice. Notably, upon CDNF treatment, *Atf4* mRNA was decreased to WT levels (**Fig. 6c**). In accordance with previous results ^10,11,35^, at 17 weeks of age the PERK, IRE-1α and ATF6 pathways were all upregulated in SOD1-G93A mice, and in particular the mRNA levels of *Chop*, which has a prominent role in apoptosis, increased more than 4-fold. CDNF treatment was able to attenuate markers from all three UPR branches (**Fig. 6b-f**). Furthermore, fluorescence staining revealed that the levels of the ER chaperone GRP78 and the phosphorylation of eIF2α were increased in lumbar MNs of 17 week-old SOD1-G93A mice, but these were efficiently reduced by CDNF administration, resulting in expression levels comparable to WT mice (**Fig. 6g-j**). SOD1 is a cytosolic protein, but mutant SOD1 has also been observed in the ER lumen ^11^. Therefore, we investigated whether CDNF treatment would affect the levels of mutant SOD1 protein and its clearance using its conformation-specific antibody B8H10 (MM070, Medimabs, Canada): we found a modest reduction in the amount of mutant SOD1 in the SOD1-G93A mice at 17 weeks (**Fig. S7**).

**Figure 6.**
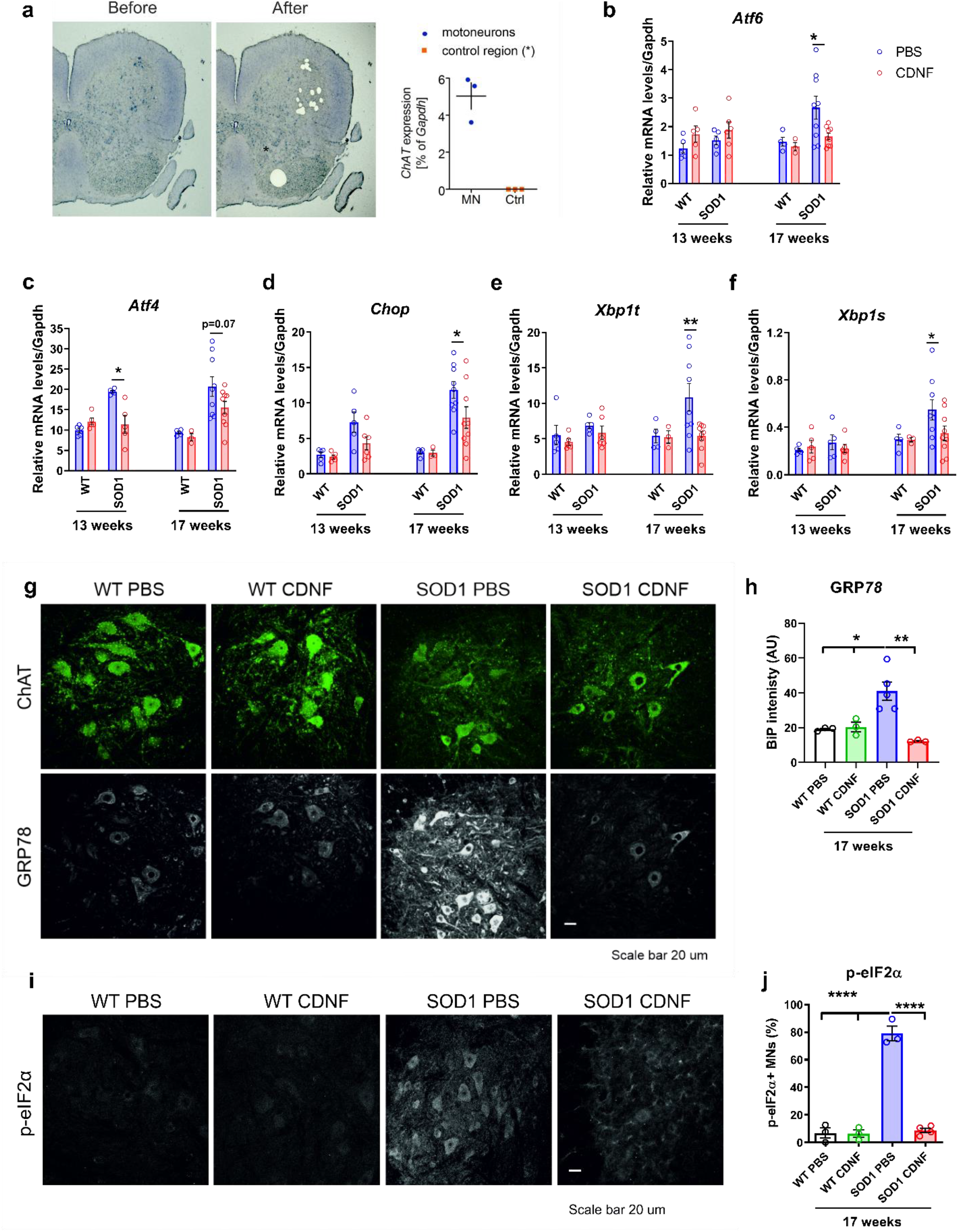
A single i.c.v. injection of CDNF decreases the expression of UPR markers in the lumbar spinal cord of SOD1-G93A animals at 17 weeks. (**a**) Representative pictures of a lumbar spinal cord section before and after the MNs dissection and mRNA level of *ChAT*, marker of spinal cord MNs, in the putative MNs area and in a control region where no MNs should be present (marked with *). (**b-f**) mRNA expression of UPR markers *Atf6, Atf4, Chop, Xbp1t* and *Xbp1s* in microdissected lumbar MNs after CDNF/PBS injection in 13 and 17 weeks SOD1-G93A mice and WT littermates. (**g**) Expression of GRP78 protein in lumbar MNs after CDNF/PBS treatment in 17 weeks SOD1-G93A mice and WT littermates. (**h**) Quantification of **g**. (**i**) Expression of p-eIF2α protein in lumbar MNs after CDNF/PBS treatment in 17 weeks SOD1-G93A mice and WT littermates. (**j**) Quantification of **i**. Mean ± SEM, n=6-9/group in **b-f**, n=3-5/group in **g-j**. *p<0.05, **p<0.01, ****p<0.0001, two-way ANOVA followed by Bonferroni post-hoc test in **b-f**, one-way ANOVA followed by Tukey post-hoc test in **h** and **j**.

### Endogenous CDNF levels change with disease progression in SOD1-G93A mice

We next evaluated the endogenous levels of CDNF in non-treated WT and SOD1-G93A mice at different disease stages, from the pre-symptomatic stage of 1 month to the endpoint of approximately 5 months, and collected lumbar spinal cord, motor cortex, and skeletal gastrocnemius muscle. We found that levels of CDNF change during SOD1-G93A lifespan. CDNF was upregulated at the pre-symptomatic stage in the spinal cord and later in the motor cortex. On the contrary, CDNF levels in skeletal muscle were significantly decreased at symptom onset stage and onwards (2, 3 and 5 months of age, **Fig. 7**), which might correlate with reduced activity of paralyzed muscle.

**Figure 7.**
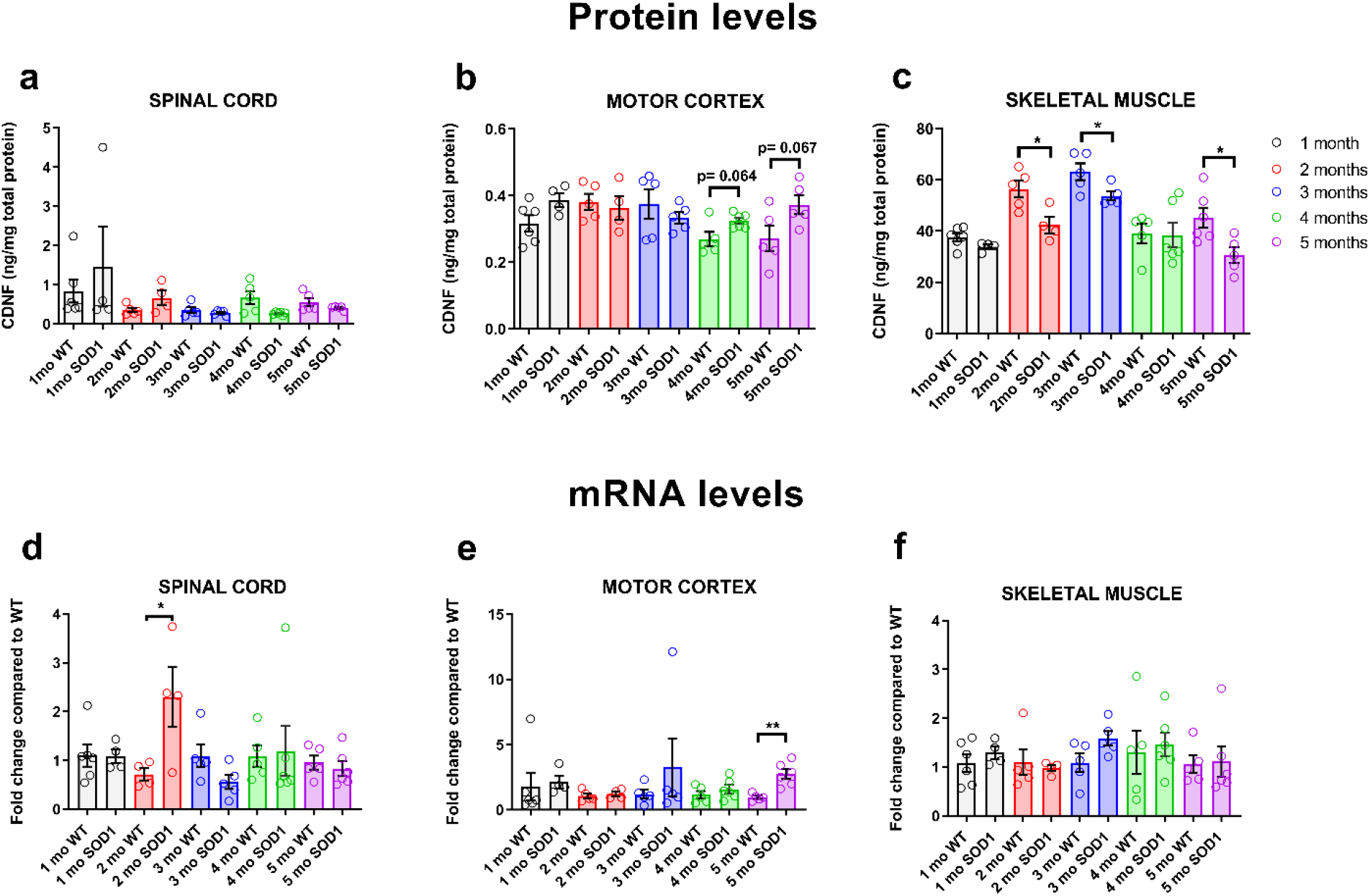
CDNF levels change with disease progression in SOD1-G93A mice. Protein and mRNA levels of CDNF were measured from the spinal cord, motor cortex and skeletal gastrocnemius muscle of SOD1-G93A and WT littermates at different time points (1,2,3,4 and 5 months of age). Protein levels of CDNF in the spinal cord (**a**), motor cortex (**b**) and skeletal muscle (**c**). In the same order, mRNA data are presented in **d-f**. Mean + SEM; n= 4-6 animals/group; * p<0.05, **<0.01, unpaired t-test.

### CDNF reduces the UPR response in a slow progressive TDP43-M337V mouse model

Since enhanced sensitivity to ER stress in SOD1-G93A MNs is detectable as from embryonic stages (**Fig.5b**) and long before the appearance of first clinical signs in the corresponding mouse model (**Fig. 4**), we sought to explore the protective effect of CDNF in a third and new model of ALS, which displayed a slow progressive development of the disease. Herein, TDP43-M337V is expressed as a transgene under the ubiquitin promoter, resulting in transgene levels that are comparable to the endogenous levels of TDP43 (**Fig. S8a-b, e-f**). Thus, this preclinical model mimics the balance between mutant and WT TDP43 levels as observed in patients with one mutant TDP43 allele. These mice do not show any severe clinical symptoms and premature death, but develop MNs loss at the age of about 12-15 months (**Fig. S8c-d**). This is an agreement with two recent studies, describing the phenotype of new mouse models in which the mutation in the TDP43 gene was introduced into the endogenous locus and where only a mild phenotype with late onset was observed ^36,37^. Mice aged more than one year and used in this study, show electrophysiological alterations such as decreased numbers of motor units and reduced motor performance on a rotarod when challenged by rocking movements or a combination of accelerating and rocking wheel movements (**Fig. S8g-h**). Signs of an enhanced ER stress were present long before an altered behavior was detected in this mouse model (**Fig. 8a**). To test the effect of CDNF, TDP43-M337V mice received a single i.c.v. injection of 10 μg of CDNF at 6 weeks of age and the ER stress response was analyzed at 6 months of age in the spinal cord and the motor cortex of the mice (**Fig. 8b**). An exacerbation of the UPR response was found in the transgenic mice compared to the WT littermates. As expected, this upregulation was not as strong as the one detected in the SOD1-G93A at late stage. However, treatment with CDNF had a similar effect in reducing the ER stress response, with a significant decrease in *Chop* and *Xbp1t* mRNA and an equivalent trend in most of the other genes (**Fig. 8c-h**). Moreover, the mRNA levels of the UPR markers were downregulated in the full lysate of motor cortex upon treatment with CDNF (**Fig. 8i**). These results are in line with the effects observed in the TDP43-M337V rat and in the SOD1-G93A mouse, and demonstrate that CDNF actively suppresses ER stress in ALS preclinical models independently of the disease etiology.

**Figure 8.**
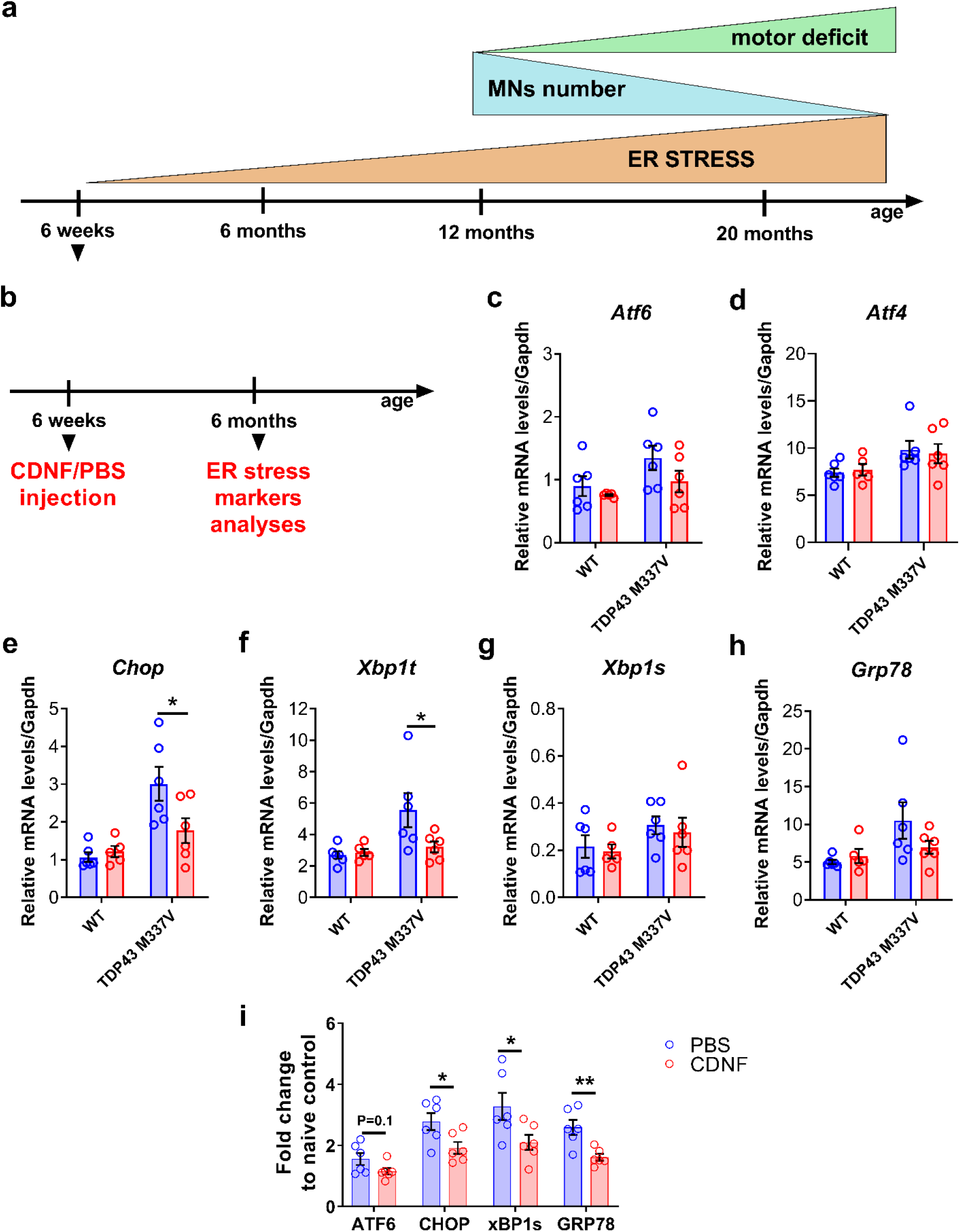
CDNF treatment decreases UPR markers expression in a novel TDP43-M337V mouse model with ER stress pathology. (**a**) Schematic representation of the appearance of ER stress and MNs loss in the novel TDP43-M337V mouse model. (**b**) TDP43-M337V mice were injected at 6 weeks of age and UPR markers were analyzed by qPCR at 6 months. (**c-h**) mRNA expression of UPR markers *Atf6, Atf4, Chop, Xbp1t, Xbp1s* and *Grp78* in microdissected lumbar MNs from 6 months TDP43-M337V mice treated with CDNF or vehicle at 6 weeks of age. (**i**) mRNA expression of UPR markers *Atf6, Chop, Xbp1s* and *Grp78* in total lysates of motor cortex from 6 months TDP43-M337V mice treated with CDNF or vehicle at 6 weeks of age; results are presented as fold change increase compared to naïve control. Mean ± SEM, n=5/6/group in **c-h**, n=5/group in **i**. *p<0.05, **p<0.01; two-way ANOVA followed by Bonferroni post-hoc test in **c-h**; unpaired t-test in **i**.

## DISCUSSION

ALS is a complex and multifactorial disease ^38,39^. Several studies have indicated that ER stress plays a pivotal role in the pathophysiology of the disease ^3–19^, thus it became an interesting target for therapeutic intervention. In this study, we explored the role of CDNF as a modulator of the ER-stress response, and show that CDNF is the first compound that can attenuate all the three UPR pathways activated by IRE-1α, PERK, and ATF6 in MNs *in vitro* and *in vivo*. This is remarkable, as previous compounds have been shown to interfere only with the PERK/p-eIF2α pathway, in particular inhibiting the de-phosphorylation of eIF2α ^10,20,21^ or by suppressing the activity of the dual leucine zipper kinase (DLK) ^40^. Small molecule allosteric inhibitors for IRE1α oligomerization, Kinase-Inhibiting RNase Attenuators (KIRA) have also been described: KIRA6 is a specific IRE-1α kinase inhibitor, whereas KIRA8 blocks IRE-1α RNase activity. They have been shown to reduce beta-cell ER stress and death, and thus ameliorate diabetes in mouse models ^41,42^. We have previously discussed that all three pathways are chronically active in ALS and that the UPR is a finely intertwined network.

Thus, targeting only one of the pathways is not sufficient and limits the efficacy of these compounds, whereas CDNF has a broader therapeutic scope than any single pathway-specific inhibitor. Furthermore, it was also reported that persistent translation inhibition by p-eIF2α, which is accentuated by drugs as salubrinal, mediates neurodegeneration in a model of prion disease, which is another protein misfolding disorder that shares several pathogenetic mechanisms with ALS ^43^. Treatment with guanabenz, which was shown to be protective in female SOD1-G93A mice by Jiang *et al.* ^21^, led to an opposite effect in another study, with an exacerbation of disease symptoms especially in male mice ^44^. These results suggest that guanabenz, especially if administrated systemically, may have detrimental effect because of a sustained block of protein translation, acceleration of apoptosis, and also because of possible side effect due to the activity of the drug as an agonist of the α2-adrenergic receptor ^44^. On the other hand, a complete restoration of protein translation by PERK inhibitor GSK2606414, albeit being neuroprotective in a prion model, was found to be highly toxic for pancreatic cells, where UPR activation is essential for physiological activity ^45^. These reports suggest the need of a drug candidate with the ability to attenuate the whole UPR network in affected neurons, without side effects. Interestingly, we found that CDNF has very little effect *in vitro* on healthy MNs but has a strong protective effect on injured and ER stressed-MNs. Broad toxicity assays for CDNF have been already carried out in non-human primates and CDNF was successfully tested in phase I-II clinical trial in Parkinson’s disease (PD) patients ^28^.

Chronically activated UPR pathways eventually lead to neuronal dysfunction and finally to CHOP-mediated cell death ^46^. CDNF significantly reduced the levels of CHOP and we postulate that CDNF inhibits CHOP-mediated apoptosis, therefore enhancing MNs survival. In addition, CDNF was shown to activate the PI3K-Akt pathway in dopamine neurons ^30^, and it warrants to be studied whether this occurs also in MNs. Importantly, the long-lasting downregulation of UPR markers in the SOD1-G93A and TDP43 mouse models by CDNF, even after a single injection of CDNF, indicates that the protective effect is maintained for a long period of time, highlighting its potential therapeutic value.

Due to the genetically variegated and multifactorial nature of ALS, creating an animal model that would properly mimic all the pathophysiological events involved in the development and progression of the disease has revealed as complex task and has led to the unsuccessful translation of drugs from the bench to the bedside ^47^. Therefore, being aware of the advantages and disadvantages of available pre-clinical model, we sought to investigate the therapeutic effect of CDNF in three rodent models with different etiology and disease development. Albeit their differences, all three models exhibit signs of ER stress and death of MNs and treatment with CDNF was able to attenuate the ER stress response and rescue MNs, irrespective of the disease etiology.

Administration of CDNF via the i.c.v. route improved the motor behavior and increased the number of surviving MNs in both the rat TDP43-M337V and in the mouse SOD1-G93A model. To the best of our knowledge, no previous drug candidate has been reported to have a protective effect in this severe and fast progressive TDP43 rat model. In a recent report, it was shown that riluzole had no therapeutic efficacy on the behavioral deficits, nor on the neuropathological features in this model ^48^. Furthermore, CDNF is the first agent showing a beneficial effect in the SOD1-G93A mice when treated at 3 months of age, i.e. after symptom onset. This is a clinically relevant time point for initiation of therapy in patients, as usually they are not diagnosed before disease onset. SOD1-G93A mice treated daily with riluzole in drinking water, starting approximately 40 days before symptom onset, showed improved survival by 13 days ^49^, whereas no increase in survival was observed after edaravone intraperitoneal administration ^50^. Additionally, the evidence from our novel TDP43-M337V mouse model that ER stress is already present before the appearance of clinical symptoms suggests that CDNF is also a strong candidate for the prevention of disease initiation in ALS cases with familial monogenic origin.

Finally, in this study we demonstrate that endogenous levels of CDNF change during disease progression, being upregulated at pre-symptomatic stages and decreasing at later stages. In particular, the decreased levels of CDNF in muscle compared to WT might correlate with paralysis and reduced activity of non-functional muscle. Furthermore, the decrease in the endogenous levels of CDNF with disease progression suggests that the administration of exogenous protein at this stage should have a therapeutic effect, which is supported by the results described here.

In conclusion, CDNF rescues MNs *in vitro* and *in vivo* by regulating the ER stress response, a key player in the pathophysiology of ALS. Hence, we propose that CDNF is a promising ER stress regulator for injured MNs that warrants further study as a drug candidate in ALS.

## MATERIALS AND METHODS

### Study design

The aim of this study was to investigate the therapeutic effect of CDNF in preclinical models of ALS and to elucidate CDNF mechanism of action in relation to ER stress. To this end, being aware of the drawbacks of available ALS pre-clinical models, we extended our research to three rodent models, which exhibited different disease development and etiology: (i) the conditional ChAT-tTA/TRE-hTDP43-M337V rat model described by *Huang et al.* ^29^, (ii) the SOD1-G93A mouse model, and (iii) a slow-progressive TDP43-M337V mouse model. The first two model were used to evaluate the effect of CDNF treatment on motor behavior and disease progression. All three models were implicated in the analysis of the ability of CDNF to attenuate the UPR at different disease stages and to rescue MNs from ER stress-induced cell death. To evaluate the ER stress response specifically in MNs, which are the most affected cell-type in ALS, we utilized three main methods: (i) primary culture of MNs derived from E13 embryos, (ii) immunohistochemical analyses of spinal cord sections with ChAT as spinal MNs marker, and (iii) qPCR analyses of lumbar MNs isolated via laser microdissection. All the specific methods and characteristics of the animal models are described below.

The experimental design for each animal models is illustrated in the main text and in the **figures 1a, 4a, and 8b**, respectively for ChAT-tTA/TRE-hTDP43-M337V rats, SOD1-G93A mice and TDP43-M337V mice.

All animal experiments were carried out following European Community guidelines for the use of experimental animals and approved by the Finnish National Experiment Board (License numbers: ESAVI/7907/04.10.07/2014 and ESAVI/8897/04.10.07/2017).

To ensure randomization in the animal studies, subjects were assigned to each treatment group on the base of their motor performance, as assessed by baseline rotarod testing. All behavior experiments and analyses were performed blindly by the investigators.

### Animal models

#### SOD1-G93A mouse model

Female and male SOD1-G93A mice (strain B6SJL-TgN (SOD1-G93A)1Gur) from The Jackson Laboratory were used in this study. SOD1-G93A express the human mutant (glycine at codon 93 to alanine) superoxide dismutase 1 gene, driven by the human SOD1 promoter. Transgenic expression was assessed by DNA tail test and qPCR (SOD1-G93A fwd: 5’-CAC GTG GGC TCC AGC ATT-3’; SOD1-G93A rev: 5’-TCA CCA GTC ATT TCT GCC TTT G-3’; internal CTR fwd: 5’-GGG AAG CTG TTG TCC CAA G-3’; internal CTR rev: 5’-CAA GGG GAG GTA AAA GAG AGC-3’). In all experiments, WT littermates were included as controls. Mice were housed according to standard conditions, with access to food and water ad libitum and a 12 hour dark/light cycle. Animals were monitored twice per week from the pre-symptomatic stage (11 weeks of age) until the endpoint for body weight changes and symptom progression. A clinical score scale from 5 to 1 was used to determine disease progression, according to ALS Therapy Development Institute (ALSTDI) guidelines: onset of disease (score 4) was determined by observing tremors in the hind limbs and gait impairment. Animals were sacrificed at the endpoint (score 1) when unable to right themselves within 30 seconds when placed on one side.

#### ChAT-tTA/TRE-hTDP43-M337V rat model

Transgenic rats were created on a sole Sprague Dawley background as described previously ^29^. Transgenic expression was assessed by PCR of rat tail DNA with the following primers: ChAT-tTA fwd: 5’-TCCAAGGCAGAGTTGATGAC-3’; ChAT-tTA rev: 5’-TGAGTTCCAGGCAAACCAAG-3’; hTDP-43 fwd: 5’-TGCGGGAGTTCTTCTCTCAG-3’; hTDP-43 rev: 5’-AGCCACCTGGATTACCACCA-3’. In all experiments, WT littermates were included as controls. Rat were housed according to standard conditions, with access to food and water ad libitum and a 12 hour dark/light cycle. hTDP43-M337V transgene activation was suppressed by addition of doxycycline (Dox) to the drinking water (50 μg/ml, Sigma Aldrich, MI, USA). Dox was constantly given to breeding rats and to pups until genotyping analyses at one month of age; after that, only TDP43-M337V^+^ rats continued to receive Dox water, while WT animals received normal water. At 60 days of age, a partial withdrawal of Dox was applied to induce transgene activation (10 μg/ml, dilution 1:5). All experiment rats were sacrificed at 21 days after transgene activation by perfusion.

#### Novel TDP43-M337V mouse model

For generation of transgenic TDP43-M337V-expressing mice, the coding sequence of TDP43 was mutated and cloned into a derivate of the lentiviral vector FUG-W ^51^. A HA-tag was introduced at the N-terminus to monitor the transgene expression. Concentrated lentiviral vector was injected into single-cell mouse embryos, which were subsequently implanted into pseudo-pregnant females. After germline transmission, animals were backcrossed for 4 generation into a Bl6/C57J background. Mice were housed according to standard conditions, with access to food and water ad libitum and a 12 hour dark/light cycle.

### Stereotaxic surgeries

#### CDNF and ^125^I-labeled CDNF

Recombinant human CDNF (rhCDNF) was produced by baculoviral expression in Sf9 insect cells ^23^ and purified by Biovian Inc. (Turku, Finland). An additional thrombin site was introduced for the removal of FLAG-6His tags. The biological activity of rhCDNF was assessed in a superior cervical ganglion neuronal survival assay ^52^. For ^125^I-labeled CDNF, CDNF (1.0 μg) was iodinated with ^125^I-Na using the lactoperoxidase method. CDNF was dissolved in 30 μl of 0.25 M phosphate buffer, pH 7.5, and mixed with ^125^I-Na (1 mCi=37 mBq; GE Healthcare). The reaction was started by adding lactoperoxidase (10μl of 50μg/ml) and 5μl of 0.05% H_2_O_2_. The mixture was incubated at room temperature for 20 min and the reaction was stopped by adding 3 vol of 0.1 M phosphate buffer, pH 7.5, containing 0.1 M NaI, 0.42 M NaCl, and 25 μl of 2.5% BSA. Free iodine and iodinated growth factor were separated on Sephadex G-25 columns (PD10; GE Healthcare). For column equilibrium and elution, 0.1 M phosphate buffer, pH 7.5, with 1% BSA was used. The iodinated growth factors were concentrated by using YM-10 Centricon columns (Thermo Scientific, MA, USA). The specific activity of ^125^I-labeled CDNF was 10^8^ cpm/μg.

#### Single injection paradigm

13 weeks SOD1-G93A mice and 6 weeks TDP43-M337V mice received 10 μg rhCDNF or vehicle (PBS) through unilateral stereotaxic injection in the brain ventricle under isoflurane anesthesia (coordinates relative to bregma and dura: A/P +0,3; L/M −1; D/V −2).

For the diffusion study, 10 μg of rhCDNF was injected unilaterally in the brain ventricle using the same brain coordinates as above. Mice were transcardially perfused at 2, or 6 hours after infusion, and brains were processed for CDNF immunohistochemistry. For the ^125^I-labeled CDNF study, five microliters of radiolabeled protein (2.5 ng) plus vehicle were injected directly into the ventricle using the stereotaxic coordinates described above. Transcardial perfusion was performed after 1,3, and 24 h under deep anesthesia. Portions of central nervous tissue were used for measurements in a gamma counter, and the remainder was sectioned for autoradiography.

#### Continuous infusion paradigm

The stereotaxic surgery protocol using Alzet osmotic pumps (model 2004, Durect Corp., CA, USA) was modified from Voutilainen *et al.* ^27^. All procedures were carried out under isoflurane anesthesia. The brain infusion cannula was implanted into the cerebral ventricle using the coordinates relative to the bregma A/P −1.0; M/L -/1.5, D/V −4.0 ^53^. It was secured to the skull via the installation of three stainless streel screws and with polycarboxylate cement (Aqua-lox; VOCO, Germany). The cannula was attached linked to the osmotic pump via a 2.5 cm long catheter.

Finally, the osmotic pump was placed into a subcutaneous pocket located between the shoulder blades. Osmotic minipumps filled with hCDNF (6 μg/day) or PBS were active for 28 days. The filling, priming, and calculation of the concentration of infused protein for the Alzet minipumps were prepared according to the manufacturer’s instructions.

### Analysis of motor behavior

#### SOD1-G93A mice

Motor behavior of animals was assessed using a battery of different tests. <u>*Accelerating rotarod:*</u> mice were tested for their ability to maintain balance on a rotating rod with accelerating speed (4-40 rpm) and the latency to fall was measured (cut off time 4 minutes; Rota-Rod, Ugo Basile, Italy). <u>*Multiple static rods:*</u> this is a series of five 60 cm long wooden rods of decreasing diameters (rod 1 - 27 mm; rod 2 - 21 mm; rod 3 - 15 mm; rod 4 - 11 mm; and rod 5 - 8 mm), each perpendicularly screwed at one end to a supporting beam. Mice are placed on the distal end of the rod, facing away from the supporting beam. The latencies to orient 180° to face the fixed end of the rod (orientation time), and then to travel 60 cm to the supporting beam (travel time), were recorded. <u>*Open field test:*</u> the locomotor and rear activity of the mice was measured for 1 hour in an open field device (Activity Monitor 5.1, MED Associates Inc., GA, USA). <u>*Grip strength:*</u> strength of forelimbs and hind limbs was measured separately every week using Chatillon force measurement device (FL, USA). The measurement of strength (Newton) was normalized on the weight of each animals (grams).

#### ChAT-tTA/TRE-hTDP43-M337V rats

To investigate the motor deficits, the rats were assessed using the accelerating rotarod test. Before the stereotaxic surgery, the rats were trained one time per day for three days, with acceleration speed (4-40 rpm) and a cutoff time of 240 seconds. Upon the baseline training, the animals were divided in balanced groups according to the measurement of the latency to fall from the rod. After transgene activation, the rats were tested three times per weeks until day 21.

#### TDP43-M337V mice

To analyze the motor performance, mice were tested for their ability to maintain balance on a rotating rod using three different setups. For the accelerating rotarod, the speed of the rod was accelerated from 10-40 rpm within 30 s. For the rocking rotarod, the speed was kept constant at 15 rpm for 35 s and then the direction of the rod turned. During the third setup, the rod was accelerated from 15 - 33 rpm in 10 sec, slowed down from 33 −15 rpm in 10 sec, and then the direction of the rod turned and the procedure was repeated.

### Immunohistochemistry

#### SOD1-G93A mice

For immunohistochemical analyses, animals were perfused transcardially with PBS and then with 4% PFA; spinal cords and brains were collected and post-fixed overnight in 4% PFA.

##### ChAT and CDNF

Thirty-micrometer-thick cryosections were cut through the spinal cord (L3-L5) and brain. Spinal cord sections were incubated with goat anti-ChAT (1:1000, Merk Millipore, MA, USA) and brain sections were incubated with rabbit anti-CDNF (1:500, ProSci, Poway, CA, USA) antibodies followed by incubation with biotinylated secondary antibodies, rabbit anti-goat or goat anti-rabbit (1:200, Vector laboratories Inc., Burlingame, USA). Antibodies were combined with standard ABC reagents (Vectastain Elite ABC kit, Vector laboratories Inc., Burlingame, USA) and the signal was visualized with 3,3’-diaminobenzidine (DAB) (Sk-4100). The area of CDNF distribution in the brain was determined by measuring the area of CDNF-immunopositive staining in square millimeters in every sixth brain section. For the diffusion of CDNF, total protein diffusion volume was estimated with the Cavalier Estimator function of the Stereo Investigator platform using a grid spacing of 500 μm as described previously ^54^. For the quantification of ChAT^+^ neurons, the number of MNs was counted from serial cryosections of lumbar region segments of the spinal cord. Every 10^th^ section was analyzed and the mean of counts on both sides of the spinal cord was statistically evaluated.

##### GRP78 andB8H10

For immunofluorescence staining, sections were kept for 2 hours in PBS solution containing 0.05% Triton X-100 and 10% normal donkey serum (NDS). The primary antibodies goat anti-ChAT (1:250, Millipore, AB114P), mouse anti-B8H10 (1:250, Medimabs), rabbit anti-SOD1 (1:500, Enzo Lifesciences, ADI-SOD-100) rabbit anti-BiP (GRP78) (1:250, Abcam, ab21685) were applied in PBS, 3% NDS, 0.05% Triton X-100, and incubated for 2 days at 4 °C. Sections were then washed with PBS (3 x 10 min) and incubated 2 hours with the appropriate secondary antibodies (Invitrogen, CA, USA) at room temperature. The sections were then rinsed again with PBS and coverslipped with Dako fluorescence mounting medium.

##### p-eIF2α

For immunohistochemistry staining of p-eIF2α (rabbit anti-p-eIF2α, 1:50, Cell Signaling, 3597L) positive MNs, 10 μm paraffin sections were deparaffinized. Heat-mediated antigen retrieval was performed using 10mM sodium citrate buffer, pH 6. The endogenous peroxidases were inactivated followed by 1 hour of blocking in TBS-T (tris-buffered saline + 0.1% tween 20) containing 10% horse serum (HS). The sections were incubated overnight with the primary antibody at 4° C. Sections were then washed with TBS-T and the proper secondary antibody was applied for 1 hour at room temperature. The sections were then incubated for 30 minutes with Avidin-biotin complex (ABC) solution, washed 3 times (3 x 10 min) with TBS-T buffer before the 3’-diaminobenzidine (DAB) reaction. The glass slides were then dehydrated with ascending concentration of ethanol and coverslipped.

Confocal images were acquired using an Olympus Fluoview 1000-BX61 microscope (Olympus, Tokyo) fitted with a 20X, 40X air or 60X immersion oil objective. The same confocal settings were maintained throughout the acquisition between groups in order to measure the intensity of expression of GRP78 and B8H10. Bright-field microscopy images were acquired with a Nikon Eclipse Ti-E microscope equipped with a 10X air objective, in order to count the motor neurons positive for p-eIF2α. Images were analyzed with FiJi, signal intensity values for the antigen of interest were calculated over 10 consecutive lumbar spinal cord Z-stack spaced 0.5 μm, after background subtraction from every different channel the intensity of GRP78 and B8H10 of ChAT positive MNs was analyzed.

#### ChAT-tTA/TRE-hTDP43-M337V rats

After PFA transcardial perfusion, spinal cord was processed and embedded in paraffin.

##### ChAT

10 μm thick paraffin sections were deparaffinized. Antigen retrieval was performed heating up the sections in the antigen retrieval solution (10mM sodium citrate buffer, pH6.0, 0.05% Tween-20). Sections were incubated with 0.3% H_2_O_2_ in TBS for 30min to block the activity of endogenous peroxidase. Sections were rinsed with TBS-T (0.1% Tween-20 in TBS) 2 times for 10 min and block with 1.5% horse serum (S-2000, Vector Laboratories) in TBS-T for 1 hour at room temperature. Sections were incubated with primary antibodies goat-anti-ChAT (1:500, Millipore, AB144P) in 1.5% horse serum over night at 4 °C. The next day, section were rinsed with TBS-T and incubated with biotinylated-anti-goat IgG (BA-9500, 1:400, Vector Laboratories) for 1 hour at room temperature. Sections were rinsed with TBS-T, incubated with avidin-biotin complex solution (PK-4000, Vector Laboratories) for 1 hour and the signal was visualized with 3,3’-diaminobenzidine (DAB) (Sk-4100). Stained sections were scanned using 3DHISTECH at the Institute of Biotechnology, HiLIFE, University of Helsinki.

##### GRP78 and p-PERK

For immunofluorescences staining, the sections underwent deparaffinization and antigen retrieval. After blocking with 1.5% donkey serum in TBS (Sigma Aldrich, MI, USA) for 1 hour at room temperature, the sections were incubated with primary antibodies goat-anti-ChAT (1:200, Millipore, AB144P), rabbit-anti-GRP78 (1:500, Abcam, ab21685), and rabbit-anti-p-PERK (Thr981, 1:200, Santa Cruz, sc-32577) in blocking solution overnight at 4 °C. The following day, the sections were rinsed with TBS-T and incubated with Alexa 488-conjugated anti-goat IgG (1:500, Thermo Fisher Scientific) and Alexa 589 conjugated anti-rabbit IgG (1:500, Thermo Fisher Scientific). Fluorescence images were acquired using confocal microscopy (LSM700, Carl Zeiss).

The sections stained with ChAT were analysed as described above. Fluorescence images were captured with 25x magnification. The intensity (pixel value) for each antigen was quantified from 4 images of the ventral horn area for each animal by Image J software (Image J, Loci at the University of Wisconsin-Madison, USA). Image J co-localization plugin was used to quantify the level of co-localization between the protein of interest and ChAT, and the results were normalized by the number of ChAT+ neurons ^55^.

#### Novel TDP43-M337V mice

After removal of the spinal cord, the tissue was post-fixed in 4% PFA for 2 h and subsequently embedded in 5% agarose. Forty micrometers thick free-floating sections were cut on a Leica 9000 s sliding microtome as collected in 0.1 M phosphate buffer (PB), pH 7.4. After incubation of the sections for 1 h in blocking solution (10% normal goat or donkey serum and 0.3% Triton X-100 in TBS-T) at room temperature, the sections were incubated for 48h at 4 °C with primary antibodies in washing solution (1% normal goat or donkey serum and 0.03% Triton X-100 in TBS-T). After three washing steps for 10 minutes in washing buffer at room temperature, the sections were incubated with fluorescently labeled secondary antibodies (diluted in TBS-T), washed again and finally mounted with Flour Save (Merck). The following primary antibodies were used: ChAT (1:1000; Millipore, MAB144P), anti-HA (1:1,000; Sigma-Aldrich, clone 3F10). Alexa-647-, Alexa-488-, and Alexa-546-conjugated secondary antibodies were from Jackson Immuno-Research Laboratories.

##### Nissl staining

Paraffin serial sections of the lumbar spinal cord were Nissl stained, and MNs were quantified on every 10^th^ section.

### Analyses of radioactive ^*125*^I-CDNF

#### Autoradiographic analysis of the distribution of ^125^I-CDNF

Mice receiving an i.c.v. injection of ^125^I-CDNF (5—10 ng) were perfused for 24 h after injection. Coronal paraffin sections (7 μm thick) were exposed to autoradiography film (Kodak Biomax MS) for 4 weeks.

#### Quantification of ^125^I-CDNF in tissues after intraventricular injection

Brain, spinal cord, and other organs were dissected and the wet tissue weighed. The amount of intraventricularly administered protein in different structures was determined at 1, 3, and 24 hours after perfusions using Perkin Elmer Wallac Wizard 1470 Gamma Counter.

### Motor neuron culture

Murine embryonic spinal MNs were isolated and cultured as described ^56^. Briefly, after dissection of the ventrolateral part of E12.5 embryos, spinal cord tissue was incubated for 15 min in 0.1% trypsin in Hank’s balanced salt solution. Cells were triturated and incubated in Neurobasal medium (Invitrogen, CA, USA), supplemented with 1× Glutamax (Invitrogen, CA, USA) on Nunclon plates (Nunc) pre-coated with antibodies against the p75 NGF receptor (MLR2, the kind gift of Robert Rush, Flinders University, Adelaide, Australia) for 45 min. Plates were washed three times with Neurobasal medium, and the remaining MNs were recovered from the plate using depolarization solution (0.8% NaCl, 35 mM KCl and 2 mM CaCl_2_) and collected in full medium (2% horse serum, 1x B27 in Neurobasal medium with 1x Glutamax). For survival assays, MNs were isolated from single embryos and 1,000 cells per cm^2^ were plated on poly-ornithine/laminin-coated dishes. MNs were counted 4 h after plating to determine the total number of plated cells. Cells were counted again at the indicated times to determine the number of surviving cells. For the survival assay 10 ng/ml of CDNF, BDNF, and CNTF were used. To analyze the protective effect of CDNF against thapsigargin-induced ER-stress, the total number of cells was determined before the addition of thapsigargin and the number of surviving cells was analyzed 32 hours after treatment. For qPCR- and Western blot analysis, embryos were genotyped during motor neuron isolation, and cells isolated from embryos with a similar genotype were pooled obtain sufficient cell numbers. For the measurement of UPR-markers, cells were treated for 8h with 5 nM thapsigargin. The concentration of CDNF was 50 ng/ml.

### Plasmid construction and lentiviral vector production

FUW-EGFP-ATF6 was generated by digestion of the FUW plasmid by *XbaI*, and the resulting 10 kB fragment was purified by gel extraction. pEGFP-ATF6 was digested by *NheI* and *XbaI*, and the resulting insert was purified by gel extraction and cloned into the FUW backbone by ligation. The *5’XbaI* site was destroyed during cloning. pEGFP-ATF6 was a gift from Ron Prywes (Addgene plasmid # 32955). Lentiviral vector was produced as previously describe ^57^. Briefly, HEK 293T were co-transfected with FUW-EGFP-ATF6 and packaging plasmids using Ca-phosphate. The medium was replaced 24 h after transfection and collected 24 h later. Subsequently, the viral vector was concentrated by ultracentrifugation.

### Immunocytochemistry

For immunocytochemistry, cells were grown on poly-L-ornithine (PORN)/Laminin coated-glass coverslips and fixed with phosphate-buffered PFA for 15 min at RT, washed three times for 5 min each with PBS, and blocked for 30 min with blocking buffer containing 10% donkey serum and 0.3% Triton X-100 in TBST (TBS-Tween). After three 5 min washes with TBST at room temperature (RT), cells were incubated with primary antibody overnight at 4°C, followed by three 5 min washes in TBST at RT, and then incubated with the appropriate fluorophore-conjugated secondary antibody for 1h at RT. Subsequently, cells were washed three times for 15 min each with TBST. Coverslips were mounted on glass slides with FluorSave (Merck Millipore, MA, USA). Cells were imaged using an Olympus Fluoview 1000i confocal microscope. The following primary antibodies were used: anti-Tuj1 (1:500; Neuromics, MN, USA MO15013), anti-ChAT (1:200; Merck Millipore, MA, AB144P). Alexa-647 and Alexa-546-conjugated secondary antibodies were purchased from Jackson Immuno-Research Laboratories. For visualization of F-actin Alexa-488 conjugated Phalloidin (Invitrogen, CA, USA) was applied to the cells during incubation with secondary antibody.

### Laser microdissection

Spinal cord samples were cryo-cut into 20 μm sections at −18°C and directly transferred onto membrane-covered slides. After 30 s of fixation in 70% Ethanol, slides where dried for 10 min at 40°C. Sections were stained for 3 s in 0.02% toluidine blue, rinsed in DEPC-treated water and decolorized for about 3 min in 75% ethanol. All solutions were prepared with DEPC-treated water. After 10 minutes of drying at 40°C, slides were stored on ice for laser microdissection, which was performed with the Leica Microsystems laser microdissection DM 6000 B. MNs and control areas from 32 spinal cord sections per mouse were selected and 4.7 μl of reverse transcriptase - lysis buffer was added and samples were stored on ice. Finally, after 2 min at 72°C, 1 min on ice and a short spin-down, 0.5 μl of reverse transcriptase were added to the MN containing samples. Reverse transcription was performed at 37°C for 2 hr, followed by a 5 min cooking step at 95°C. Samples were stored at −20°C until processing for qPCR. For qPCR samples were diluted 1:10 and two 2 μl were used as a template.

### Quantitative RT-PCR

#### For animal tissue

Total RNA was isolated from the lumbar spinal cord, motor cortex, and gastrocnemius muscle using TriReagent (Sigma Aldrich, MI, USA) according to the manufacturer’s instructions. To ensure the removal of any DNA residue, RNA samples were treated with DNA-free kit (Thermo Scientific, MA, USA). cDNA was synthesized from 1 μg RNA using RevertAid Premium Reverse Transcriptase (Thermo Scientific, MA, USA) and Oligo(dT)18) primers, according to the manufacturer’s protocol.

#### For MNs

Total RNA was extracted from cultured MNs using the RNAeasy RNA Isolation Kit (Qiagen, Germany) following the manufacturer’s instruction. At least 100 ng of total RNA was reverse transcribed using a First Strand cDNA Synthesis Kit (Thermo Scientific, MA, USA). The cDNA was diluted 1:10 and 1 μl was subsequently used as a template for qPCR.

Expression of UPR marker and CDNF mRNAs was quantified using the LightCycler® 480 Real-Time PCR System and Lightcycler 480 SYBR Green I master mix (Roche, Switzerland). β-actin was chosen as a reference gene. Results were obtained with the Basic Relative Quantification Analysis of the LightCycler® 480 Real-Time PCR System software (Roche, Switzerland) and analyzed using the ΔΔCt method. The following primers for *CDNF* were used: CDNF fwd 5’-GTGAGGCCATTGGTCAAACT; CDNF rev 5’-CGTGCTGTGCAATGCTTAAT; and primers for *Chop, ATF6a, Grp78, xBP1s, xBP1s, ATF4*, and *actin* as previously described ^58^.

### Western blot analyses

#### For animal tissues

For Western blot analysis of the lumbar spinal cord, the tissue was dissected, flash-frozen, and homogenized in lysis buffer (10 mM Tris, pH 7.5, 150 mM sodium chloride, 1% NP40, 0.5 mM sodium deoxycholate) supplemented with protease- and phosphatase inhibitors (Roche, Switzerland) by mechanical homogenization on ice. Homogenized tissues were incubated for 45 min on ice, and vortexed every 15 min. After incubation, the samples were centrifuged at 21.000 x g for 15 min at 4°C. Subsequently, the supernatant was collected and the protein concentration was determined by Bradford Protein Assay (Bio-Rad). Equal amounts of protein were separated by SDS-PAGE, and transferred to PVDF or nitrocellulose membranes. Membranes were blocked in TBST containing 5% milk powder for 1 h at RT, probed with primary antibodies overnight at 4 °C, and incubated with horseradish peroxidase-conjugated secondary antibodies for 1 h at RT.

The following primary antibodies were used: anti-HA (1:8,000; Sigma-Aldrich, clone 3F10), anti-TDP43 (1:4,000; Proteintech, 10782-2-AP), and anti-Tuj1 (1:4,000; Neuromics, CH23005).

#### For MNs

Cultured motor neurons were lysed in 2x Laemmli buffer, boiled for 5 minutes and sonicated. Equal amounts of protein were separated by 10% SDS-PAGE and transferred to PVDF or nitrocellulose membranes (Pall, NY, USA) by wet blotting for 1h (120V, 4°C). Membranes were blocked in TBST with 5% milk powder or 5% BSA for at least 1 h at RT, probed with primary antibody overnight at 4 °C, and incubated with horseradish peroxidase-conjugated secondary antibody for 1 h at RT. The following primary antibodies were used: anti-phospho-eIF2α (1:1000, 3597, Cell Signaling, MA, USA), rabbit anti-eIF2α (1:1000, 5324, Cell Signaling, MA, USA) anti-Chop (1:2000, 2895, Cell Signaling, MA, USA), anti-Tuj1 (1:8000, MO15013, Neuromics, MN, USA), Protein expression was quantified using Image J 1.47v software (Image J, Loci at the University of Wisconsin-Madison, USA) and the results are expressed as relative density compared to the loading control.

### Enzyme-linked immunosorbent assay (ELISAs)

The levels of endogenous CDNF protein in skeletal muscle, spinal cord, and motor cortex from 1 to 5 month old SOD1-G93A and WT mice were analyzed using in-house-built mouse CDNF ELISA ^59^. Shortly, a MaxiSorp (Nunc) immunosorbent 96-well plate was coated overnight at +4°C with 1 μg/ml of goat anti-mouse CDNF (R&D Systems, AF5187) in coating buffer (15mM sodium carbonate, 35mM sodium bicarbonate, pH 9.6). The next day, the plate was blocked with blocking buffer (3% BSA (Merck Millipore, MA, USA 820451) in PBS) for 2h at room temperature. The standard curve (recombinant mouse CDNF, R&D Systems, 5187-CD) and the study samples, diluted in blocking buffer, were applied to the plate in duplicate and incubated over night at +4°C with agitation. Bound CDNF was measured by antibody detection (rabbit anti-CDNF 0.2 μg/ml; *12*), incubated for 3 h at +37°C, followed by horse radish peroxidase (HRP)-linked donkey anti-rabbit IgG (1:2,000; GE Healthcare, NA9340V) for 2h at RT. For the color reaction, plates were incubated with 3,3’,5,5’-tetramethylbenzidine according to manufacturer instructions (DuoSetELISA Development System, R&D Systems) and absorbance was read at 450 nm (VICTOR^3^ spectrophotometer, Perkin Elmer). All antibodies and samples were applied to the plate in 100 μl volumes, while washing and blocking steps were performed in 200 μl volumes. Between steps, wells were washed repeatedly with 200 μl of washing buffer (0.05% Tween 20 in PBS).

Sensitivity of the mouse CDNF ELISA was 6 pg/ml, as determined by the mean response of ten blank samples added with three standard deviations (SD). The assay dynamic range was 15.6 to 500 pg/ml and mean intra- and inter-assay precision 10.3 and 6.8 %CV, respectively. Within the assay dynamic range, the individual back-calculated accuracy values were ±20% relative error (%RE = derived concentration / expected concentration x 100%) and precision max 15% coefficient of variation (%CV = SD / mean x 100%). Intra-assay precision was determined by measuring three samples with varying concentrations of recombinant mouse CDNF in replicates of ten on different parts of a plate, while inter-assay precision was determined by running three samples with different recombinant CDNF concentrations in duplicate in a minimum of five independent assays. *Cdnf^-/-^* ^60^ mouse tissue lysates were negative on CDNF ELISA, implicating high specificity to the analyte.

The amount of recombinant human CDNF was quantified from motor cortex and spinal cord 1 and 2 hours after i.c.v. injection of 10 μg CDNF. The quantitation was done by in-house-built human CDNF ELISA, previously described in detail ^61^ and used to quantitate human CDNF in rodent samples ^62,63^. Respective tissue from vehicle-injected mice were used as negative controls.

### Statistical analysis

Statistical analysis was performed using the IBM SPSS Statistic 22 and GraphPad Prism 8.0. *Kaplan-Meier and the log-rank test* were used for survival comparisons. *Mann-Whitney test* was used to compare the clinical status of SOD1-G93A animals. *Unpaired Student t-test* was used for multirod analyses, to analyse CDNF levels, and to compare the levels of mutated SOD1 protein in spinal cord sections. *One way ANOVA* followed by Tukey post-hoc test was used to compare the levels of ChAT, GRP78, p-PERK and p-eIF2α in immunohistochemical analyses. *Two-way ANOVA* followed by Bonferroni or Sidak post-hoc test was used for embryonic motor neuron survival, qPCR and WB analyses. *Repeated measures ANOVA* followed by Tukey post-hoc test was used for rotarod and grip strength analyses.

All results are expressed as mean ± SEM and all statistical tests are two-sided.

## SUPPLEMENTARY MATERIALS

Fig. S1. Levels of TDP43 proteins in thapsigargin and tunicamycin stressed motor neurons are unchanged after CDNF treatment.

Fig. S2. Distribution of CDNF and ^125^I-CDNF after a single intracerebroventricular injection.

Fig. S3. CDNF reaches the MNs in the spinal cord and motor cortex.

Fig. S4. Single intracerebroventricular injection of CDNF ameliorates SOD1-G93A symptoms and increases limbs strength.

Fig. S5. Cultured motor neurons derived from WT and SOD1-G93A mice treated with BDNF and CNTF or CDNF show no differences in morphology.

Fig. S6. CDNF reduces mRNA levels of UPR markers in treated motor neurons.

Fig. S7. Effect of CDNF on mutated SOD1 levels.

Fig. S8. Characterization of novel TDP43 M337V mouse model of ALS.

## Acknowledgments

Katrina Albert, Emilia Hella, Pushpa Khanal, Ella Montonen, Anni Saukkonen, Matilda Sinkko and Janika Sirjala are thanked for their technical help.

