## Supplementary material for "CDNF rescues motor neurons in three animal models of ALS by targeting ER stress": Suplementary materials and figures

### SUPPLEMENTARY MATERIALS

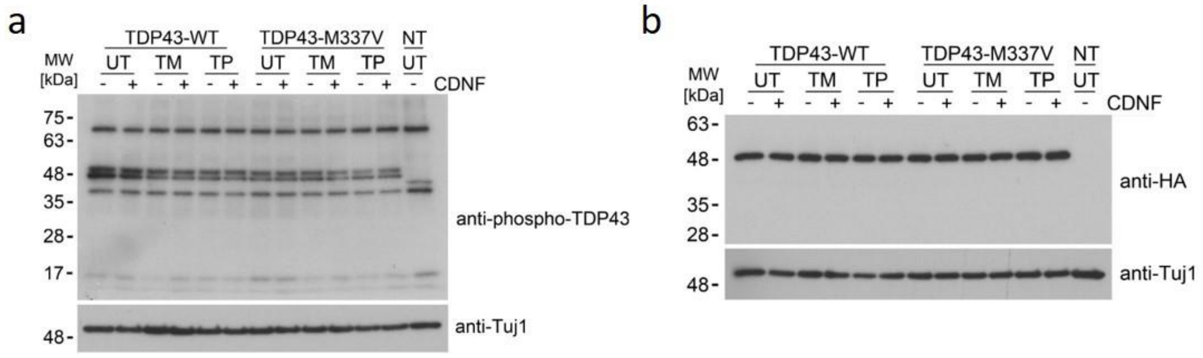


**Figure S1 | Levels of TDP43 proteins in thapsigargin and tunicamycin stressed motor neurons are unchanged after CDNF treatment.** Representative western blot pictures of p-TDP43 (**a**) and total TDP43 protein (**b**, tagged with HA).

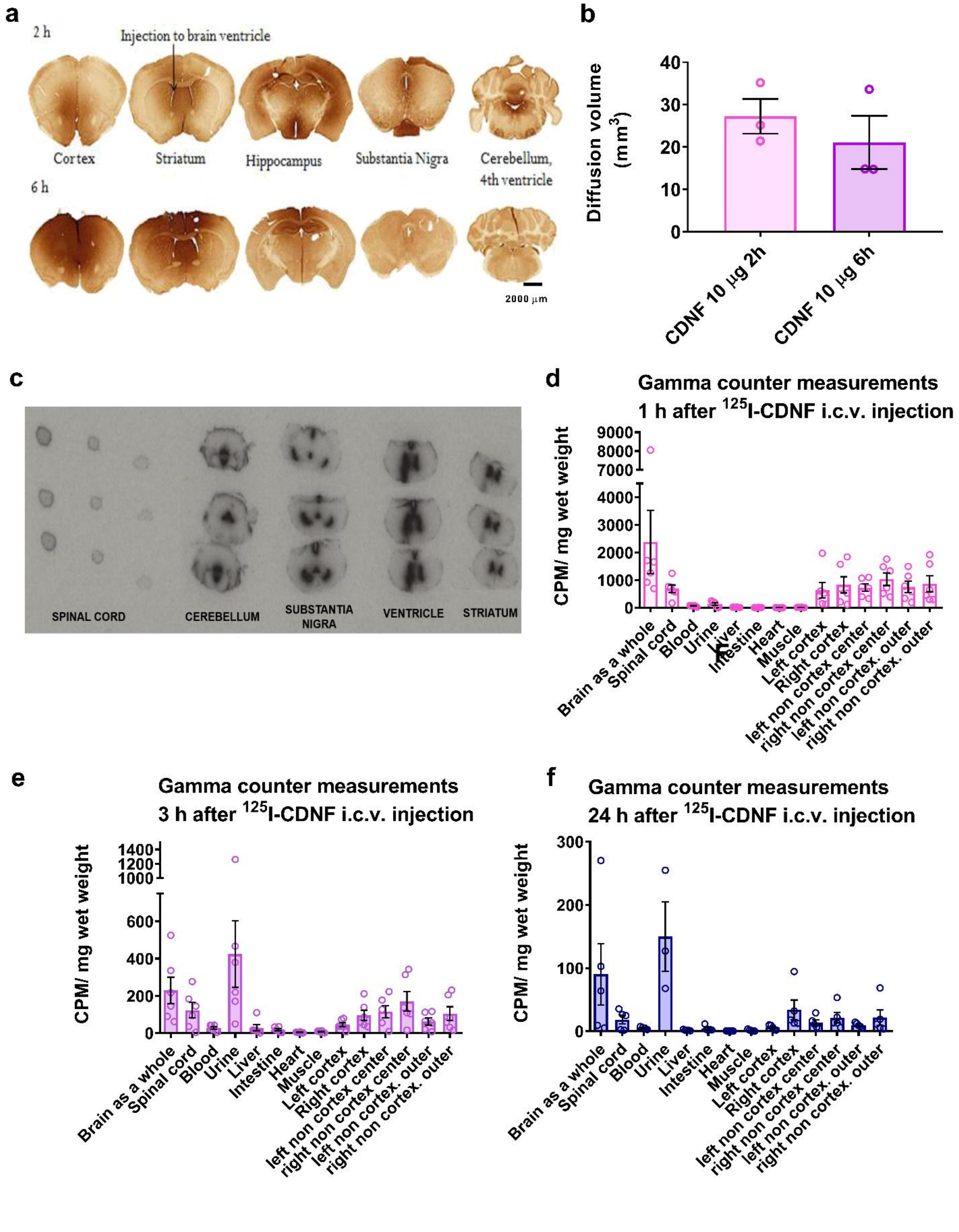


**Figure S2 | Distribution of CDNF and $^{125}$I-CDNF after a single intracerebroventricular injection**. (a) Representative immunohistochemical staining for CDNF in a coronal section through the brain two and six hours after i.c.v injection of rhCDNF. (**b**) Diffusion volume of rhCDNF protein through the brain parenchyma 2 hours and 6 hours after i.c.v. injection. **c**, Photomicrograph of a representative autoradiographic film of $^{125}$I-CDNF distribution 1 hour following i.c.v. injection. **d-f**, Quantification of $^{125}$I-CDNF in different organs respectively 1, 3 and 24 hours after i.c.v. injection. Mean ± SEM of 3 animals/group in **b** and of 6 animals/group in **d-f.**

**a**

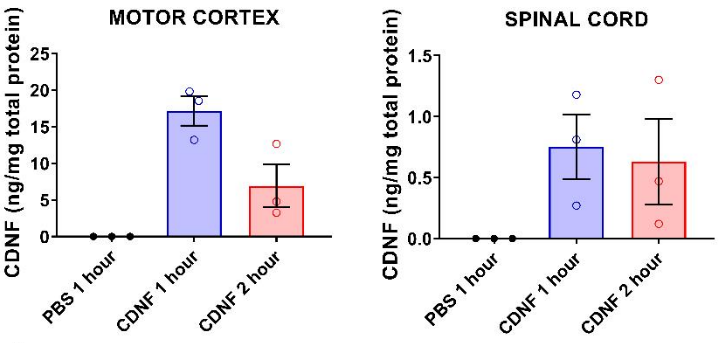


**b**

| | % of 10 µg injected i.c.v. | |
|---|---|---|
| | MOTOR CORTEX | SPINAL CORD |
| 1 hour after the injection | 0.28% ± 0.04 | 0.008% ± 0.003 |
| 2 hours after the injection | 0.091% ± 0.03 | 0.007% ± 0.006 |

**c**

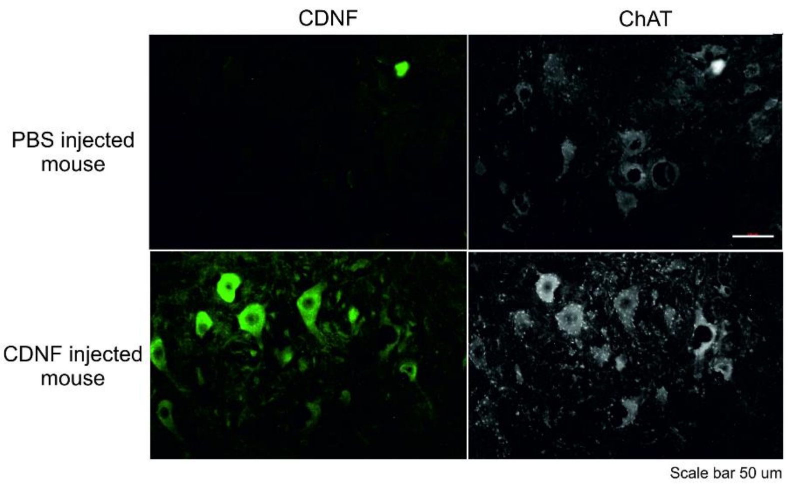


**Figure S3 | CDNF reaches the MNs in the spinal cord and motor cortex.** (**a**) ELISA quantification of the amount of CDNF that reaches the spinal cord and motor cortex after i.c.v. injection of 10 µg of CDNF. 3 animals were analyzed per group: 3 mice injected with PBS, 3 mice injected with CDNF and sacrificed 1 hour later; 3 mice injected with CDNF and sacrificed 2 hours later. (**b**) Percentage of total CDNF injected that reached the spinal cord and motor cortex 1 and 2 hours after the i.c.v. injection. (**c**) Representative pictures of fluorescence immunostaining of lumbar spinal cord sections of mice injected with CDNF or PBS.

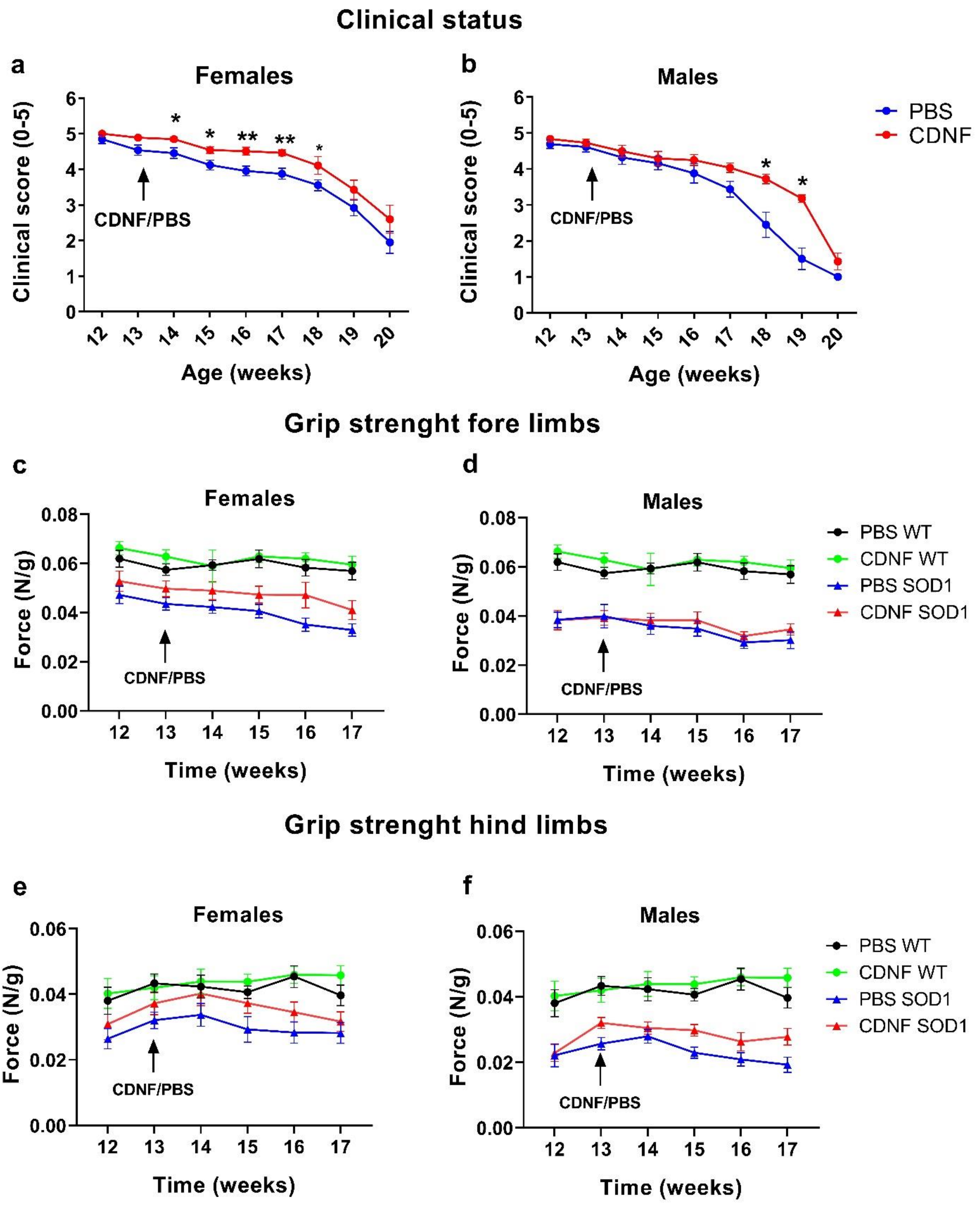


**Figure S4 | Single intracerebroventricular injection of CDNF ameliorates SOD1-G93A symptoms and increases limbs strength.** (**a-b**) Development of symptoms in CDNF and PBS-treated SOD1-G93A female and male mice measured with ALS Therapy Development Institute (ALSTDI) scoring scale. (**c-f**) Measure of limbs strength with a grip strength device in females and males in fore limbs (**c-d**) and hindlimbs (**e-f**); strength results (in Newton) are normalized on the weight of each mouse (in grams). Mean ± SEM, n=18-20 for females and 14-15 for males in **a-b**, n=15 in females and 10 in males in **c-f**; * $p<0.05$; **$p<0.001$ Mann-Withney test in **a-b**; repeated measure ANOVA followed by Tukey post-hoc test for **c-f**.

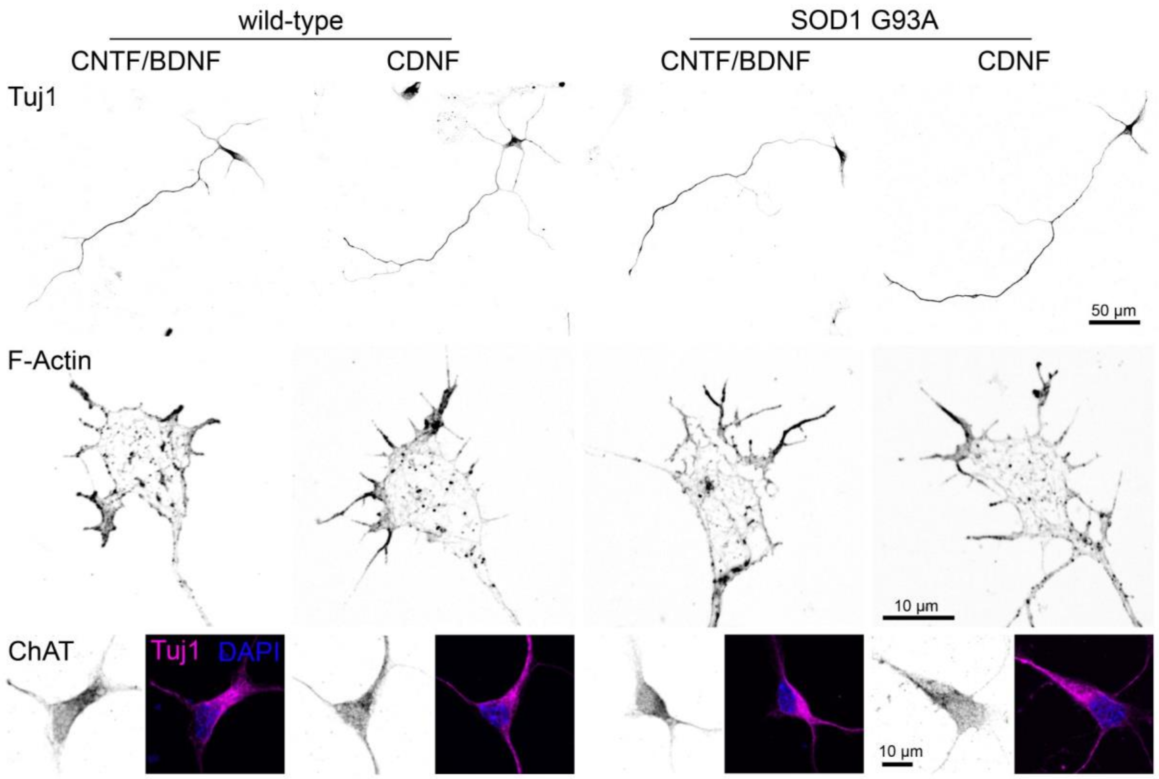


**Figure S5 | Cultured motor neurons derived from WT and SOD1-G93A mice treated with BDNF and CNTF or CDNF show no differences in morphology.** Representative images of motor neurons stained for Tuj1. For visualization of the growth cone, MNs were stained with Alexa-488 conjugated Phallodin. Immunohistochemical stainings for the MN marker ChAT, co-stained with Tuj1 and DAPI.

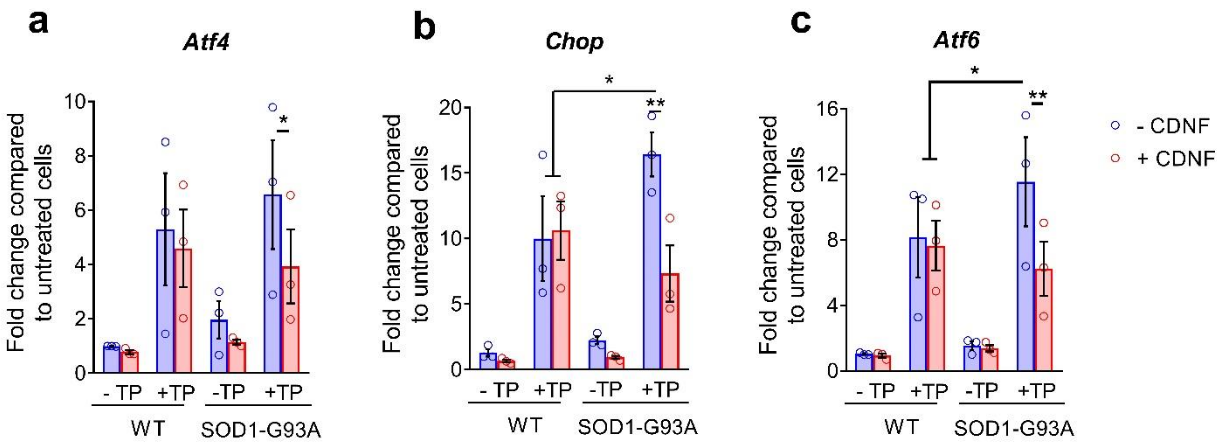


**Figure S6 | CDNF reduces mRNA levels of UPR markers in treated motor neurons.** mRNA expression of UPR markers *Atf4* (**a**), *Chop* (**b**) and *Atf6* (**c**) in thapsigargin-stressed MNs in absence or with CDNF treatment. Mean ± SEM of 3 independent experiments; *<0.05; **<0.01, two-way ANOVA followed by Bonferroni post-hoc test.

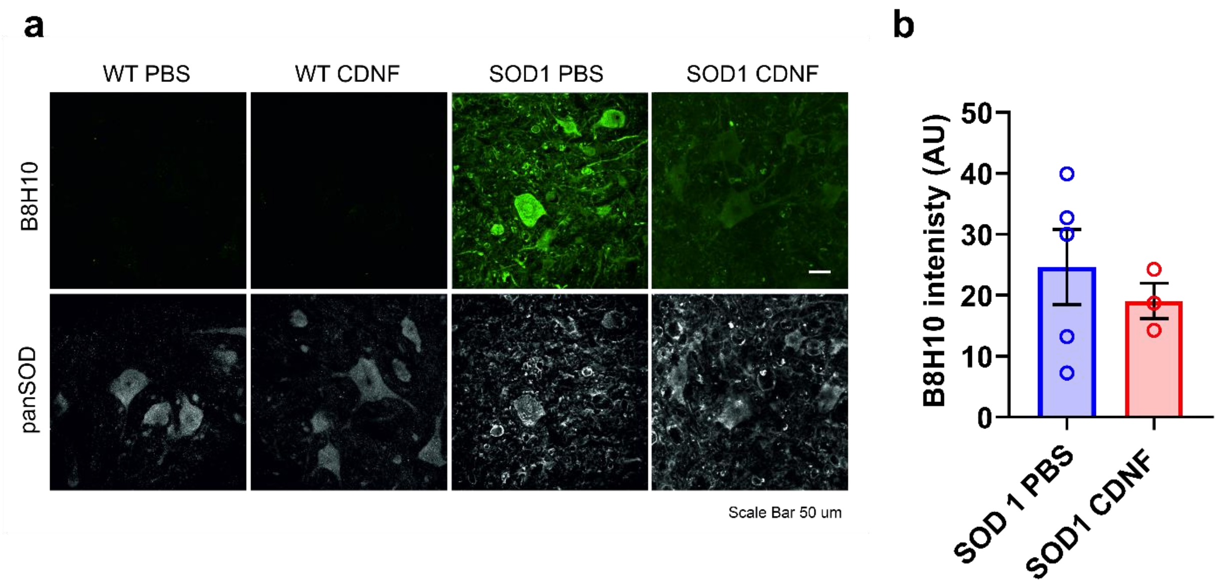


**Figure S7 | Effect of CDNF on mutated SOD1 levels.** (**a**) Representative pictures of mutated (B8H10) and total SOD1 protein immunostainings in lumbar section of CDNF/PBS-treated SOD1-G93A mice and WT littermates. (**b**) Quantification of B8H10 intensity. Mean ± SEM, n=3-5. Unpaired t-test

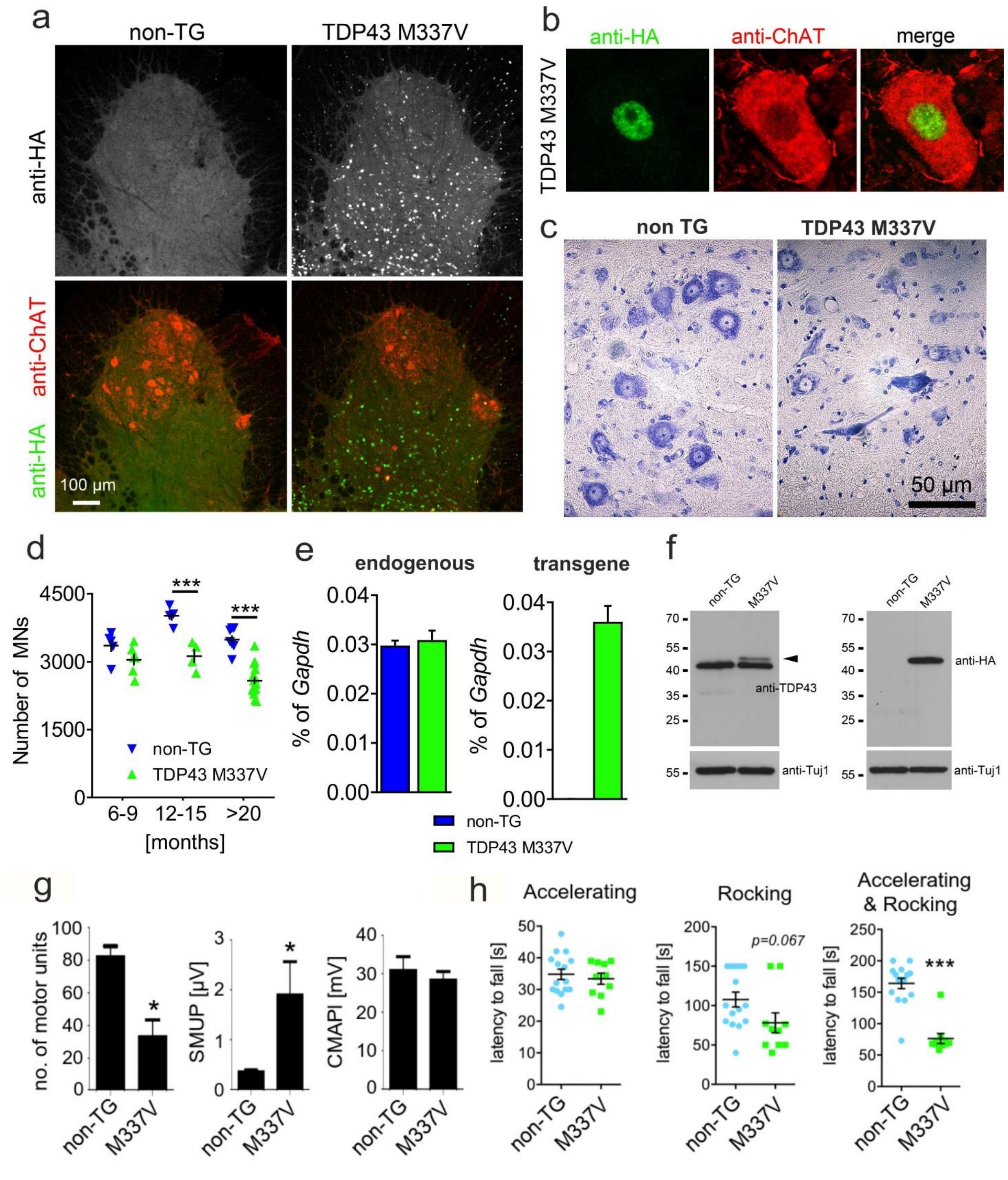


**Figure S8 | Characterization of novel TDP43 M337V mouse model of ALS.** (**a-b**) Spinal cord cross-sections from control and HA-tagged TDP43-M337V expressing animals were stained against ChAT and HA to confirm the expression of the transgene. (**c**) Nissl staining of spinal cord cross-sections from control and TDP43-M337V expressing animals. (**d**) Quantification of motor neurons'

number,*** $P < 0.001$ at 12-15 and >20 months. (**e**) mRNA levels of endogenous TDP43 and mutant TDP43 M337V analyzed by qPCR. (**f**) Western blot analysis of endogenous and transgenic TDP43 M337V. The arrowhead points to the additional band in transgenic animals detected by the anti-TDP43 antibody. (**g**) Electrophysiological measurements. The number of motor units, the single muscle unit potentials (SMUP), and the compound muscle action potentials were analyzed (CMAP), *$P < 0.05$ for motor units and SMUPs; (**h**) The motor performance of control and transgenic animals were analyzed by Rotarod analysis. The following conditions were used for the analysis. Accelerating: 10-40 rpm in 30 sec; Rocking: 15 rpm turn after 35 sec; Accelerating & Rocking: 15-33 rpm in 10 sec, 33-15 rpm in 10 sec turn, ***$P < 0.001$. TG=transgenic.
